# *In vivo* ElectroChromic Shift measurements of photosynthetic activity in far-red absorbing cyanobacteria

**DOI:** 10.1101/2024.06.01.595941

**Authors:** Julien Sellés, Jean Alric, A. William Rutherford, Geoffry A. Davis, Stefania Viola

## Abstract

Some cyanobacteria can do photosynthesis using not only visible but also far-red light that is unused by most other oxygenic photoautotrophs because of its lower energy content. These species have a modified photosynthetic apparatus containing red-shifted pigments. The incorporation of red-shifted pigments decreases the photochemical efficiency of photosystem I and, especially, photosystem II, and it might affect the distribution of excitation energy between the two photosystems with possible consequences on the activity of the entire electron transport chain. To investigate the *in vivo* effects on photosynthetic activity of these pigment changes, we present here the adaptation of a spectroscopic method, based on a physical phenomenon called ElectroChromic Shift (ECS), to the far-red absorbing cyanobacteria *Acaryochloris marina* and *Chroococcidiopsis thermalis* PCC7203. ECS measures the electric field component of the trans-thylakoid proton motive force generated by photosynthetic electron transfer. We show that ECS can be used in these cyanobacteria to investigate *in vivo* the stoichiometry of photosystem I and photosystem II and their absorption cross-section, as well as the overall efficiency of light energy conversion into electron transport. Our results indicate that both species use visible and far-red light with similar efficiency, despite significant differences in their light absorption characteristics. ECS thus represents a new non-invasive tool to study the performance of naturally occurring far-red photosynthesis.

## 1 Introduction

In oxygenic photosynthesis, the energy of sunlight is used by photosystem II (PSII) and photosystem I (PSI) to transfer electrons from H_2_O to NADPH. The intersystem electron transport chain is composed of the thylakoid membrane located plastoquinone pool and cytochrome *b*_6_*f*, and of the lumen located plastocyanin or cytochrome *c*_6_. This flow of electrons is coupled to the generation of a trans-thylakoid proton motive force (*pmf*) that drives ATP synthesis by the ATP synthase. The NADPH and ATP are then used for the fixation of CO_2_ into carbohydrates and for other metabolic pathways requiring the input of energy and reducing power. By converting the virtually unlimited sunlight into organic matter, oxygenic photosynthesis is the main biological process that sustains the production of biomass on our planet. However, the total efficiency with which oxygenic photosynthesis converts sunlight into biomass is no higher than ∼2.5-3.5% in optimal conditions, even below the theoretical maximum estimated to be ∼5-6%, with more than 60% of losses occurring at the level of the light reactions[1].

The first factor limiting the efficiency of oxygenic photosynthesis is that it only uses visible light, and thus only a restricted portion of the solar irradiation[2]. The photosystems of most cyanobacteria and of all photosynthetic eukaryotes, all having a common ancestral origin[3,4], contain exclusively chlorophyll *a* (Chl *a*), that mostly absorbs blue (Bx and By, or Soret, absorption peak) and red (Qy absorption peak) light. PSII contains 35 Chl *a* molecules, while PSI contains ∼95 Chl *a* molecules, with the precise number depending on the organism[5]. Additionally, the soluble phycobilisome antennas of cyanobacteria (PBS, containing bilins) and the transmembrane light harvesting complexes of plants and algae (LHC, containing Chl *b* or *c* and carotenoids) increase the absorption of green-orange light[6].

Once photons are absorbed, their energy is converted into a pigment’s excited state. The excitation energy then migrates from the antenna pigments to the reaction centre of the photosystems, where specialised chlorophylls (primary electron donors) use it to do photochemistry, *i.e.* charge separation. The photochemical trapping of excitation energy at the level of the reaction centres is in competition with its dissipation at the level of the antenna pigments. The primary electron donors of PSII and PSI are therefore red-shifted compared to most antenna chlorophylls, to promote energetically downhill transfer of excitation to the reaction centre. At physiological temperature, though, excitation can also be transferred energetically uphill. As a consequence, since the primary electron donor in PSII (absorbing at ∼680 nm) is less red-shifted compared to the bulk antenna chlorophylls than that in PSI (absorbing at ∼700 nm), the photochemical trapping efficiency of excitation energy is lower in PSII (∼85% [7]) than in PSI (>95% [8]). This limits the quantum yield of charge separation, and thus the proportion of absorbed light energy that is stored in the cell. It has been proposed that the reason why the primary electron donors in PSII and PSI cannot use longer wavelength light to do photochemistry is that the energy contained in red photons (∼1.82 eV for PSII and ∼1.77 eV for PSI) is the minimal amount required for promoting forward electron transfer and stabilising charge separation. This is especially the case for PSII, that needs to extract electrons from the highly stable H_2_O molecules [9–11]. Therefore, most of the oxygenic photoautotrophs do not do photochemistry using lower-energy far-red light, despite its high abundance in the solar emission spectrum.

Some oxygenic photoautotrophs can marginally extend their light use into the far-red through their light harvesting antenna proteins causing a red shift of a small number of Chl *a* molecules that can transfer the excitation energy uphill to the reaction centre. Such red-shifted Chl *a* molecules are mostly found in the core antenna of PSI in cyanobacteria[8,12] and in PSI-specific LHCs in eukaryotes[13]. Incorporating red-shifted Chl *a* in the antenna of PSII would further decrease its photochemical trapping efficiency, increasing radiative dissipation of excitation energy. Red-shifted LHCs have been found associated with PSII only in a few algae[14–16], in some of which they have indeed been shown to drastically decrease the photochemical quantum yield[14,15]. In most oxygenic photoautotrophs, longer light wavelengths are thus preferentially absorbed by PSI with respect to PSII. Therefore, when moving towards longer excitation wavelengths, charge separation rates decrease faster in PSII than in PSI, and electron donation to PSI becomes limiting for its turnover [17,18]. This excitation imbalance between the photosystems increases the losses in the conversion of absorbed light into electron transport along the intersystem chain, which are estimated to be >17% already under optimal conditions with balanced excitation[17].

Remarkably, despite all these energetic and kinetic constraints, some cyanobacteria have evolved the ability to do oxygenic photosynthesis using far-red light. The first to be discovered was *Acaryochloris marina* (*A. marina*), an epiphytic species that lives in marine environments enriched in far-red light, and mostly shaded from visible light[19,20]. The photosystems of *A. marina* contain only a few Chl *a* molecules, with the others being replaced by the far-red absorbing chlorophyll *d* (Chl *d*). These include the primary electron donors in both Chl *d*-PSII and Chl *d*-PSI (absorbing at ∼720 nm and ∼740 nm, respectively)[21]. *A. marina* also possesses phycobilisomes that absorb visible light (PBS-Vis), and transmembrane Chl *d*-containing CP43-like antennas that are associated to PSII[22,23]. Later, other cyanobacteria species were discovered that contain canonical Chl *a* photosystems and PBS-Vis when grown in visible light, but replace them with red-shifted variants when grown in environments enriched in far-red light[24,25]. In this process, canonical photosystems are replaced with isoforms that bind ∼90% Chl *a* and ∼10% of the far-red absorbing Chl *f*[11,25], plus one Chl *d* in the PSII of some species[11,26]. One of these far-red chlorophylls constitutes the charge-separating pigment in Chl *f*-PSII (absorbing at ∼720 nm)[11,27], while the identity of the primary electron donor in the Chl *f*-PSI is still debated[11,28–30]. In far-red light, in addition to the visible-absorbing phycobilisomes, these cyanobacteria also express far-red absorbing variants of the allophycocyanin subunit (PBS-FR)[25].

Since the far-red part of the solar spectrum is for the most part unused by chlorophyll *a*-based oxygenic photoautotrophs, Chl *d*- and *f*-containing cyanobacteria have a competitive advantage in conditions where visible light is scarce, as testified by their widespread distribution[31]. Naturally occurring far-red photosynthesis is of high interest also for its potential exploitation for sustainable biotechnological applications. Its introduction into crop plants and microalgae could lead to better light usage and result in higher biomass yields, especially in conditions where shading becomes limiting for photosynthesis (*e.g.* dense canopy or microalgae bioreactors)[9,32–34]. Nonetheless, at present we have only a limited understanding of how the presence of red-shifted pigments affects the overall photosynthetic efficiency in cyanobacteria in which they are native - and thus it is difficult to predict how they may affect other organisms in which they might be engineered in the future.

In the Chl *f*-containing cyanobacteria, the presence of long-wavelength pigments in the core antenna of PSII decreases its photochemical trapping efficiency, both in isolated cores and *in vivo*[35], although this loss of efficiency seems to be partially mitigated by excitation energy transfer from PBS-FR *in vivo*[36]. In contrast, the presence of Chl *f* in the core antenna of Chl *f*-PSI has little effect on its photochemical trapping efficiency[35], similar to what happens in Chl *a*-PSI containing red-shifted Chl *a* antennas[37]. In the case of *A. marina*, excitation energy trapping in Chl *d*-PSII is also lower than in Chl *a*-PSII, although this decrease is less than in the Chl *f*-containing variants[35]. On the other hand, the Chl *d*-PSII seems to suffer from higher probabilities of charge recombination leading to the production of damaging reactive oxygen species[34]. These differences between the two types of far-red PSII could be explained in terms of differences in the pigment composition and in the redox tuning of the cofactors involved in the stabilization of the charge separated state[34]. There is therefore growing evidence (recently reviewed by Elias, Oliver and Croce[38]) that the presence of far-red pigments affects the photochemical efficiency of photosystems, especially of PSII, but their impact on the overall photosynthetic light conversion efficiency *in vivo* is still unclear. For example, it is currently unknown how the variations in pigment composition of the photosynthetic apparatus affect the distribution of excitation energy between the photosystems, and what impact this has on the overall electron transport activity in far-red absorbing cyanobacteria. The complexity of these pigment variations renders their impact on the *in vivo* photosynthetic efficiency challenging to study. In addition, the distribution of excitation energy between PSII and PSI will depend on their relative amounts, which vary considerably in cyanobacteria depending on growth conditions[39].

In photosynthetic eukaryotes, a powerful spectroscopic method based on a physical phenomenon called ElectroChromic Shift (ECS) is widely used to measure the electron transport activity *in vivo*[40]. The ECS-based method measures the changes in the absorption of photosynthetic pigments induced by the electric field component (Δψ) of the *pmf* generated across the thylakoid membrane by the photosynthetic chain. Having an amplitude that is linearly dependent on the electric field, these absorption changes constitute a direct and quantitative measurement of photosynthetic activity. ECS can be used to measure the stoichiometry of the photosystems, their maximal rates of charge separation for a given illumination (that are proportional to the size of their light harvesting antenna), and the total electron transport rates[17,40]. More recently, we adapted this method to the model Chl *a*-containing cyanobacteria *Synechocystis* sp. PCC6803 (*Synechocystis*) and *Synechococcus elongatus* PCC7942 (*S. elongatus*)[41]. We demonstrated that the ECS signals of these cyanobacteria correspond to absorption changes of carotenoids, and that their amplitude is linearly proportional to the trans-thylakoid Δψ.

Although ECS represented a new tool to study the *in vivo* electron transport activity in cyanobacteria, its use in species other than *Synechocystis* and *S. elongatus* had not been reported until now. Here we present the characterisation of the ECS signals in two far-red absorbing cyanobacteria, the Chl *d*-dominated *A. marina* and the Chl *f*-containing *Chroococcidiopsis thermalis* PCC7203 (*C. thermalis*), also illustrating how the variations in pigment composition can affect *in vivo* spectroscopy measurements. We report the use of ECS to measure the photosynthetic efficiency of the two far-red absorbing cyanobacteria as a function of the illumination wavelength, in comparison with the model Chl *a*-containing species *S. elongatus*.

## 2 Materials and Methods

### 2.1 Cyanobacteria cultures

*Synechococcus elongatus* PCC7942 (*S. elongatus*) was grown in liquid BG11 medium[42] at 25°C under constant illumination with white light (SMD3528 60LED/M, RS Components) at ∼3 W m^-2^ (∼15 μmol photons m^−2^ s^−1^), with shaking. Both *Acaryochloris marina* (*A. marina*) and *Chroococcidiopsis thermalis* PCC7203 (*C. thermalis*) were grown at 30°C under constant illumination with far-red light (750 nm, Epitex; L750-01AU) at ∼3 W m^-2^, bubbling the cultures with air. *A. marina* was grown in a modified liquid K-ESM medium containing 14 µM iron[43], *C. thermalis* was grown in liquid BG11 medium.

### 2.2 Isolation of photosystems and Clear-Native PAGE (CN-PAGE)

*A. marina* and *C. thermalis* membranes were prepared as previously described[34]. For PSI and PSII isolation, membranes were adjusted to 1 mg Chl ml^-1^ in ice-cold buffer (50 mM MES-NaOH pH 6.5, 5 mM CaCl_2_, 10 mM MgCl_2_, 1.2 M betaine) and solubilised with 1% (w/v) n-Decyl-Beta-Maltoside (Sigma) at 4°C in the dark with constant stirring for 1 hour. Non-solubilised material was removed by ultracentrifugation at 160.000x*g* at 4°C for 1 hour. Photosystems were isolated from the solubilised material using sucrose gradient ultracentrifugation followed by ion exchange chromatography, as previously described[11]. For CN-PAGE analysis, samples corresponding to 1 µg Chl were separated on a 4-12% NuPAGE Bis-Tris gel (Invitrogen, Thermo Fisher Scientific) following the manufacturer’s instructions.

### 2.3 Absorption spectra of cells and isolated photosystems

Absorption spectra of whole cells and isolated photosystems were measured between 400 and 800 nm with a Cary 50 UV-Vis spectrophotometer (Agilent). To minimise light scattering, whole cell spectra were recorded by squeezing cell pellets in a thin layer between two microscope glass slides and placing them directly in front of the spectrophotometer detector.

### 2.5 Inhibitors and mediators

3-(3,4-dichlorophenyl)-1,1-dimethylurea (DCMU, Sigma) was dissolved in ethanol and used at a final concentration of 20 µM; hydroxylamine (HA, Sigma) was dissolved in water and used at a final concentration of 1 mM; 5-methylphenazinium methyl sulphate (PMS, Sigma) was dissolved in water and used at a final concentration of 15 µM.

### 2.4 Spectroscopy setups

Three experimental setups were used in this work.

<u>A laboratory-built optical parametric oscillator (OPO) laser-based spectrophotometer</u>[41]. In this setup, short monochromatic measuring pulses (duration 5 ns, 1 nm full width at half maximum, FWHM) were provided by an OPO (Horizon, Amplitude) pumped by an Nd:YAG laser (Surelite II, Amplitude). The measuring pulses were split between two closed horizontal cuvettes, one used as a reference (measuring light only) and one to measure absorption changes induced by actinic flashes. In this setup, transient absorption changes induced by the actinic flashes were measured by subtracting the reference signal from the measurement signal. Saturating single-turnover flashes (690 nm, duration 6 ns) were provided by another Nd:YAG laser pumping an OPO (Surelite II, Amplitude). The light-detecting photodiodes were protected from transmitted and scattered actinic light and fluorescence of the samples by BG39 filters (Schott).

<u>A laboratory-built xenon flash lamp-based spectrophotometer</u>. In this setup, short measuring pulses (duration 1.5 µs) of monochromatic light (3 nm FWHM) were provided by a xenon flash lamp (20W C13316, Hamamatsu) and a high-throughput (f/2) monochromator (Jobin-Yvon). The measuring pulses were split (by a randomized fibre optics bundle) between a reference and a measurement open square cuvette, as in the OPO-based spectrophotometer. Saturating single-turnover flashes (duration 1.5 µs) were provided by another xenon flash lamp (20W C13316, Hamamatsu). For measurements in the 400 to 570 nm region, the actinic xenon flash lamp was filtered with a 665 nm long-pass filter (RG665, Schott), and the light-detecting photodiodes were protected by BG39 filters (Schott). For measurements in the 645 to 780 nm region, the actinic xenon flash lamp was filtered with a BG39 filter (Schott), while no filters were used on the light-detecting photodiodes. Actinic multi-turnover pulses and continuous illumination were provided by LEDs with emission peaks at 658 and 732 nm (PowerStar ILH-ON01-HYRE-SC211-WIR200 and ILH-ON01-FRED-SC211-WIR200, RS Components). The intensities of the actinic LED lights were measured using a Macam model Q.101 radiometer/photometer (Macam Photometrics) equipped with a radiometric adaptor. The emission spectra of the actinic LEDs were recorded using a Black-Comet UV-VIS CXR-25 spectrometer (StellarNet). Both emission intensities and spectra of the actinic LEDs were measured at the end of the optic fibre delivering the light to the measurement cuvette.

<u>A laboratory-built LED-based Joliot Type Spectrophotometer (JTS)</u>. In this setup, short measuring pulses (duration 15 µs) were provided by a broad-emission white LED (COB V6HD THRIVE WHT SQ 6500K, Bridgelux) filtered via band-pass interference filters (Edmund Optics, 10 nm FWHM). The measuring pulses were split between an open square cuvette placed in front of the measuring diode and a reference diode. In this setup, measurements in the presence of measuring light only were performed on each biological sample, and subtracted from corresponding measurements performed in the presence of actinic illumination. Single-turnover saturating flashes (700 nm, duration 6 ns) were provided by an OPO pumped by a Nd:YAG laser (Surelite II, Continuum), multi-turnover actinic light was provided by orange-red LEDs. In this setup, the light-detecting photodiodes were protected from fluorescence of the samples by BG39 filters (Schott) and from actinic laser light by LPF650 filters (CVI Laser Optics) when measuring ECS, and by RG695 filters (Schott) when measuring P_700_^+^/P_740_^+^. For fluorescence kinetics, measuring pulses were filtered using a 480 nm filter, to predominantly excite chlorophyll, and the light-detecting photodiodes were protected by a BG39 filter (Schott, reference diode) and a RG665 filter (Schott, measuring diode).

### 2.5 Flash-induced difference absorption spectra

Spectra in Fig. 3 were recorded using the OPO-based spectrophotometer. Cells were harvested, resuspended in the respective supernatants and incubated in the dark in closed cuvettes in the presence of DCMU, HA and PMS for at least 1 hour before starting the measurements. A total of 8 actinic flashes were integrated for each detection wavelength.

Spectra in Fig. 4 were recorded using the xenon-based spectrophotometer. Cells were harvested, resuspended in the respective growth medium containing 20% (w/v) ficoll and incubated for at least 1 hour in the in the presence of DCMU and HA in anoxic conditions before starting the measurements. Anoxia was established in the reference and measurement cuvettes by adding 50 mM glucose and 200 units ml^-1^ glucose oxidase to the cell suspension and overlaying the samples with mineral oil to prevent gas exchange. A total of 8 actinic flashes were integrated for each detection wavelength.

Spectra in Fig. S5 were recorded using the xenon-based spectrophotometer. Cells were harvested, resuspended in the respective growth medium containing 20% (w/v) ficoll, and added with DCMU. Cells were maintained in oxygenated conditions by regular mixing. A total of 4 actinic flashes were integrated for each detection wavelength.

### 2.6 ElectroChromic Shift measurements

Flash-dependent ECS kinetics were measured with the JTS as *ΔI/I* 505 - 480 nm for *S. elongatus* and *C. thermalis* and as *ΔI/I* 520 - 546 nm for *A. marina*. The amounts of active photosystems present in each sample were quantified as the ECS amplitude recorded at 500 µs after a saturating flash (in control conditions for PSII+PSI, and in the presence of DCMU and HA for PSI only). A total of 8 actinic flashes were integrated for each detection wavelength. Where indicated, anoxia was established in the measuring cuvette by adding glucose and glucose oxidase, as described in section 2.5.

Electron transport rates were measured using the xenon-based spectrophotometer. ECS was measured as *ΔI/I* 500 - 485 nm for *S. elongatus*, as *ΔI/I* 500 - 480 nm for *C. thermalis*, and as *ΔI/I* 520 - 546 nm for *A. marina*. Cells were harvested, resuspended in their own supernatant at the desired concentration and incubated for at least 1 hour in the respective growth conditions and then for 10 minutes in darkness before starting the measurements. The amounts of active photosystems present in each sample were quantified using a saturating flash as described above. For all electron transport rate measurements, cells were maintained in oxygenated conditions by frequent mixing. The maximal electron transport rates (*I*, in e^-^ s^-1^ PSI+PSII^-1^) were calculated based on the ECS amplitude recorded after a 200 µs illumination with the indicated LED light intensities, normalised on the PSII+PSI amounts measured by flash ECS. A total of 60 measurements were integrated for each detection wavelength, for each actinic light condition. The steady-state electron transport rates (*J*, in e^-^ s^-1^ PS ^-1^) were calculated based on the amplitude of ECS decay during 5 ms of dark intervals during continuous illumination with the indicated LED light intensities, normalised on the amounts of active photosystems measured by flash ECS. For each actinic light condition, a total of 40 measurements were integrated for each detection wavelength.

### 2.7 P_700_^+^/P_740_^+^ measurements

Flash-dependent P_700+_ kinetics in *S. elongatus* and *C. thermalis* were measured as *ΔI/I* 705 - 740 nm, and P_740+_ kinetics in *A. marina* were measured as *ΔI/I* 740 nm, using the JTS. Total P_700+_/P_740+_ amounts were measured as the *ΔI/I* induced by multi-turnover saturating light pulses (3000 µmol photons m^-2^ s^-1^, duration 800 ms) given in the presence of DCMU and HA. A total of 3 measurements were integrated for each detection wavelength.

## 3 Results and Discussion

### 3.1 Pigment composition of *A. marina* and *C. thermalis* grown in far-red light

The pigment composition of the cyanobacteria used in this study is a key determinant of how efficiently they can use light at different wavelengths. Figure 1 shows whole cell absorption spectra of *S. elongatus* grown in white light (panel A) and of *A. marina* and *C. thermalis* grown in 750 nm far-red light (panels B and C, respectively). In all species, the blue region of the spectrum is dominated by the Soret absorption of the PSII and PSI chlorophylls. The main Soret absorption peak is red-shifted in *A. marina* (∼460 nm) with respect to the other two species (∼440 nm), because in Chl *d* both the Soret and the Qy absorption peaks are red-shifted with respect to Chl *a*[6]. In the case of *C. thermalis*, the photosystems contain ∼90% Chl *a* and ∼10% Chl *f*, that is red-shifted only in the Qy region, so their absorption in the Soret is similar to that of Chl *a* photosystems. In the orange-red region, the absorption spectrum of *S. elongatus* cells presents a peak at ∼625 nm, corresponding to PBS-Vis, and one at ∼675 nm, corresponding to the Qy absorption of Chl *a* photosystems.

**Fig. 1.**
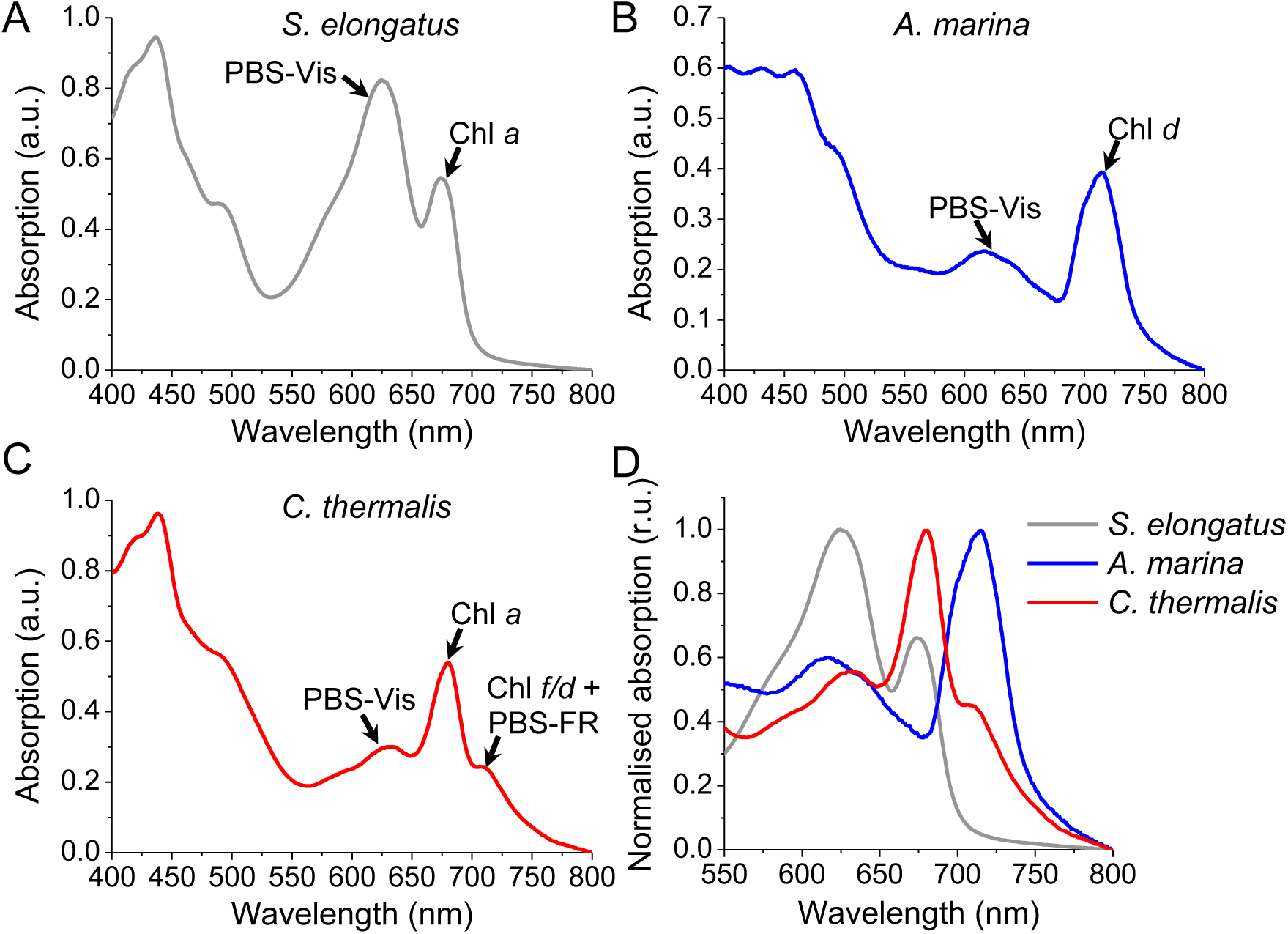
Absorption spectra of *S. elongatus* (A), *A. marina* (B) and *C. thermalis* (C) whole cells recorded between 400 and 800 nm. Black arrows indicate the peaks corresponding to visible light-absorbing phycobilisomes (PBS-Vis), far-red light-absorbing phycobilisomes (PBS-FR), and Chl *a*, Chl *d* and Chl *f* contained in photosystems. (D) Enlargement of the absorption spectra normalised to their respective maximum in the 550 nm to 800 nm region.

The absorption spectrum of *A. marina* also contains a peak corresponding to PBS-Vis, while the Qy peak of the Chl *d* photosystems is red-shifted to ∼715 nm. In *C. thermalis,* the peaks corresponding to the PBS-Vis and to the Qy absorption of the ∼90% Chl *a* present in the photosystems are also visible. An additional peak at ∼710 nm corresponds to the combined Qy absorption of the 10% Chl *f* present in the photosystems and the absorption of the far-red allophycocyanin (PBS-FR). Altogether, the absorption of *S. elongatus* cells drops drastically after 700 nm (Fig. 1D), while that of both *A. marina* and *C. thermalis* is extended in the far-red, although the two species have very different absorption profiles.

The absorption spectra of intact cells are informative but do not provide information on the actual wavelength-dependent absorption cross-section of PSI and PSII, which would be required to predict how efficiently they could use light of increasing wavelength to do photosynthesis. In *A. marina*, the absorption spectra of the isolated Chl *d-*PSI and Chl *d-*PSII are very similar, with only a minor red-shift (∼2 nm) in the Qy absorption peak of PSI (Fig. 2A and C). Therefore, in this species, PSI does not absorb light at much longer wavelengths than PSII. On the other hand, the absorption spectrum of Chl *d-*PSII contains a peak at ∼630 nm corresponding to PBS-Vis that co-purify with it, suggesting their tight association *in vivo*. PSII might therefore absorb more visible light than PSI in *A. marina*. The PBS-Vis are also visible in the CN-PAGE analysis of the purified *A. marina* fraction (Fig. 2C), together with a Chl-containing band of smaller size than PSII that could correspond to the CP43-like antenna that has been previously found to be associated with it[22]. Since this antenna also predominantly binds Chl *d*, its contribution cannot be distinguished from that of the PSII cores in the absorption spectra. No significant amounts of PBS or CP43-like antenna co-purify with *A. marina* PSI, which in our preparation migrates in the CN-PAGE as a monomer. Although published structures of *A. marina* PSI indicate that it normally occurs in trimeric form[44,45], the absence of some subunits in such structures suggests some biochemical instability that could explain the monomerization observed in our sample.

**Fig. 2.**
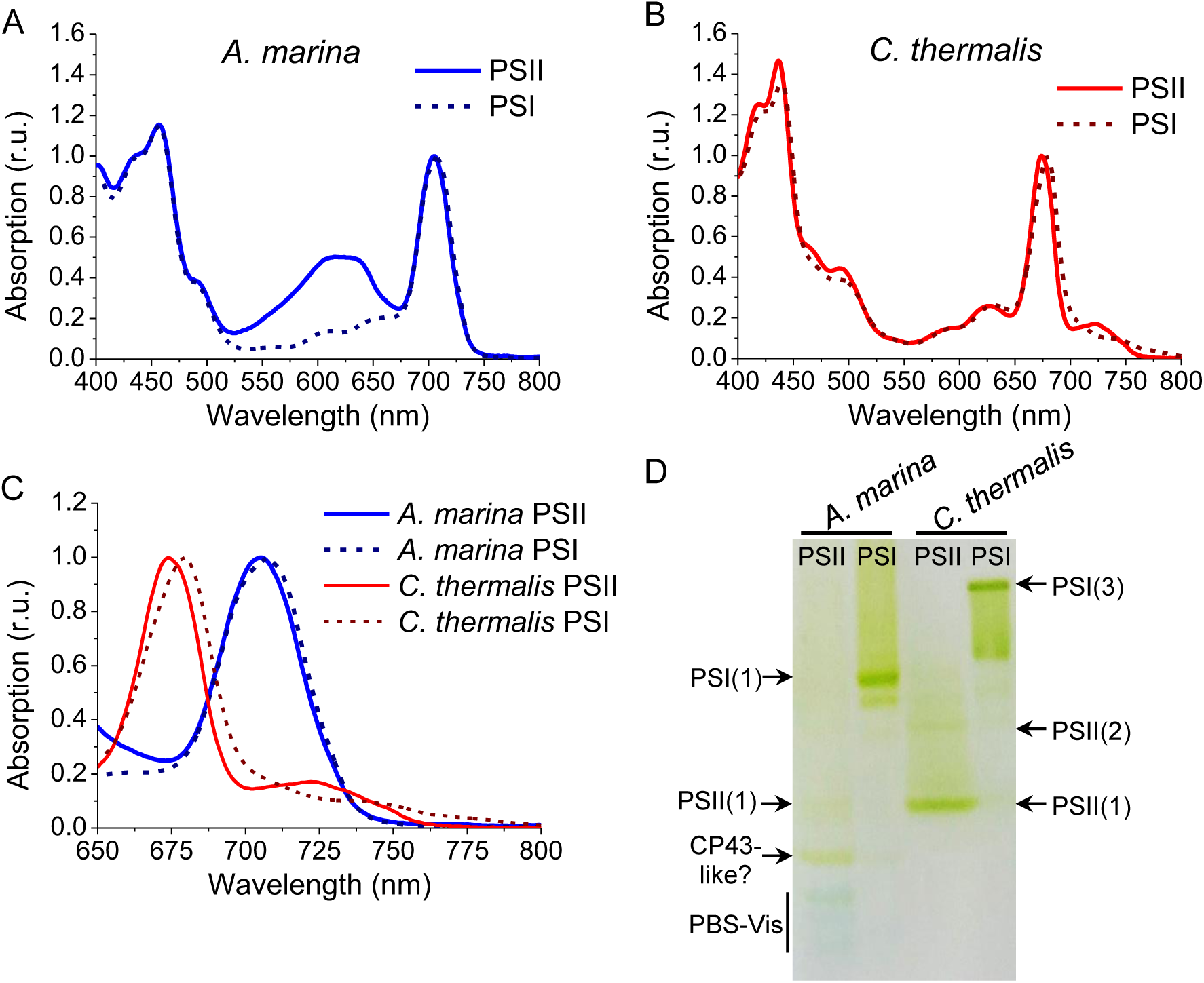
Spectral and biochemical characterisation of isolated far-red photosystems. (A) and (B) Room temperature absorption spectra of isolated PSII and PSI from *A. marina* and *C. thermalis*, respectively. All spectra are normalised to their maximal chlorophyll Qy absorption. The Qy region (650 to 800 nm) of all spectra are shown on an expanded scale in panel (C). (D) CN-PAGE of the isolated photosystems from panels A and B. Indicated are the bands assigned as PSI trimers and monomers (PSI-3 and -1), PSII dimers and monomers (PSII-2 and -1), PBS-Vis, and the band tentatively assigned as a CP43-like antenna in *A. marina*.

In the case of *C. thermalis* (Fig. 2B and C), the absorption spectra of Chl *f*-PSII and Chl *f*-PSI differ significantly in the Qy region, as previously reported[11]. The peak of the Chl *a* bulk antenna is red-shifted by ∼6 nm in Chl *f*-PSI relative to Chl *f*-PSII. The 5 far-red chlorophylls present in Chl *f*-PSII (1 Chl *d* and 4 Chl *f*[11]) absorb between ∼720 and ∼760 nm, while the 8 Chl *f* present in Chl *f*-PSI[11] extend the absorption up to ∼800 nm. Selective excitation of PSI at longer wavelengths could thus be possible in this species. No significant amounts of PBS-Vis or PBS-FR could be detected in our samples, although the far-red allophycocyanin can be found associated with both *C. thermalis* Chl *f*-PSII and Chl *f* -PSI in different biochemical preparations[11,27]. While the far-red allophycocyanin has been shown to transfer excitation to PSII in *C. thermalis* cells[36], it is unclear how much excitation is transferred to the photosystems by the PBS-Vis, that are mostly confined to the external periphery of the cytoplasm[46].

Altogether, the absorption spectra of cells and isolated photosystems indicate that in both far-red cyanobacteria excitation imbalance between the two photosystems could occur, although probably at different illumination wavelengths. The PSI of *S. elongatus* also contains a few red-shifted antenna chlorophylls that extend its absorption to ∼710 nm[47]. Despite their small absorption cross-section, they could sustain selective excitation of PSI. In the following, we will describe the characterisation of the ECS signals in *A. marina* and *C. thermalis*, and how ECS-based measurements could be used to investigate the light usage efficiency of cyanobacteria with different pigment composition, as a function of illumination wavelength.

### 3.2 Spectra and linearity of ECS signals in *A. marina* and *C. thermalis*

To characterise ECS signals in *A. marina* and *C. thermalis*, we recorded flash-induced absorption changes in the presence of the PSII inhibitors DCMU and HA and of the redox mediator PMS. PMS acts as a PSI electron donor and thus minimizes absorption changes due to the oxidation/reduction of the P_A_P_B_ chlorophylls of PSI (P_700_ in Chl *a*-PSI[48] and Chl *f*-PSI[11], and P_740_ in Chl *d*-PSI[49,50]) and of the hemes in cytochrome *b_6_f*, while maintaining ECS-related absorption changes[41].

In both far-red species, the ECS spectra obtained (Fig. 3) resemble red-shifts of carotenoids, as previously published for *S. elongatus* and *Synechocystis*[41]. The ECS spectrum of *C. thermalis* (Fig. 3B) presents the same negative peak at 480 nm and positive peak at 500 nm as in *S. elongatus*. A negative peak at 485 nm is present also in the ECS spectrum of *A. marina* (Fig. 3A), but the positive feature is broader and peaks around 515-520 nm, similar to the ECS spectrum of plants and green microalgae[40,51]. Based on the spectra, we assumed that the amplitude of ECS signals in *C. thermalis* could be measured as the difference between the light-dependent absorption changes at ∼500 and ∼480 nm, like in *S. elongatus*. It remained to be determined whether in *A. marina* ECS could be measured with this same combination of wavelengths or rather as the difference between 520 and 546 nm. This difference is often used in plants and green algae to measure ECS correcting for signals arising from the oxidation and reduction of the cytochrome *f* subunit of cyt *b*_6_*f*[52].

**Fig. 3.**
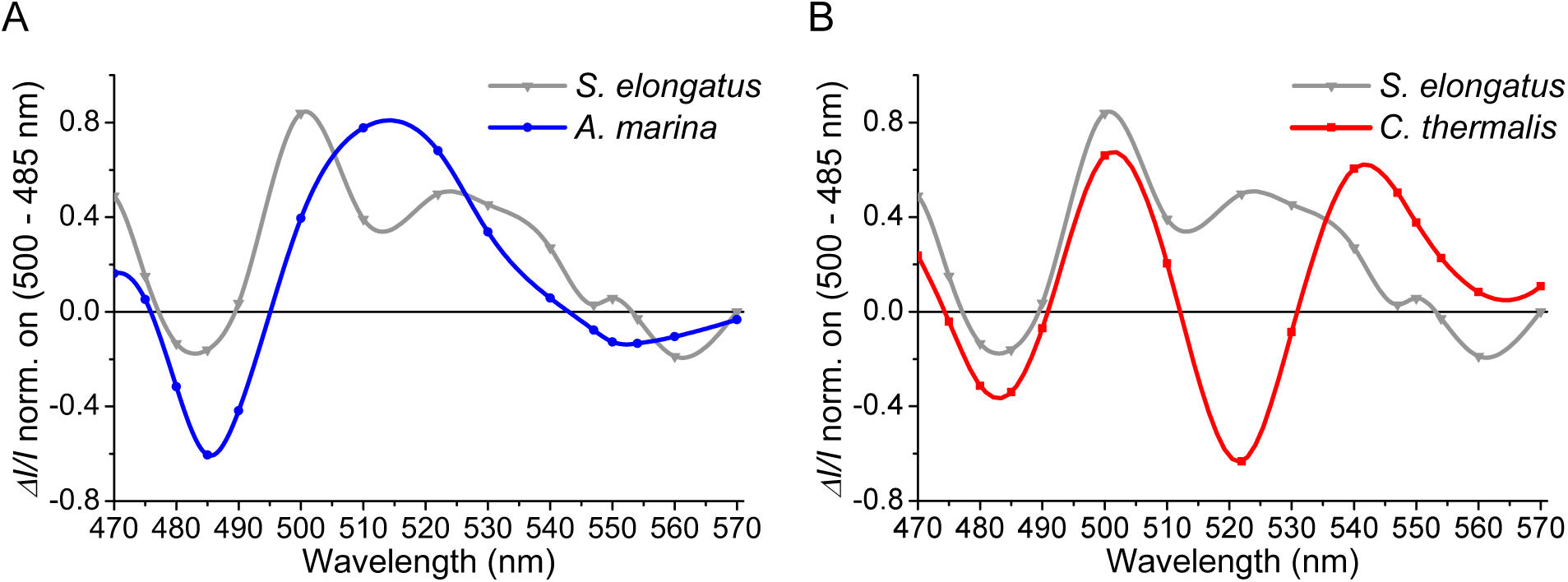
Flash-induced ECS spectra in *A. marina* and *C. thermalis*. (A) and (B) Spectra of absorption changes in *A. marina* and *C. thermalis* cells, respectively, measured at 5 ms after a single-turnover flash in the presence of DCMU, HA and PMS. The spectra are compared with those recorded in the same conditions in *S. elongatus* and previously published[41]. All spectra are normalised to the difference between the absorption changes at 500 and 485 nm.

To determine the best combination of wavelengths to measure ECS in *A. marina* and validate those for *C. thermalis*, we recorded flash-induced absorption spectra in the absence of PMS (Fig. 4). In these conditions, ECS dominates the long-lived spectral components (tens to hundreds of ms) in both species (Fig. 4C and D), while the spectra recorded at 500 µs and 20 ms after the flash also contain redox-dependent absorption changes in addition to ECS. The redox components can be better seen when subtracting the long-lived ECS from the spectra recorded at the shorter times (Fig. S1), although this subtraction is less effective in *C. thermalis* (Fig. S1B), where the ECS decays significantly between 500 µs and 200 ms, than in *A. marina* (Fig. S1A).

**Fig. 4.**
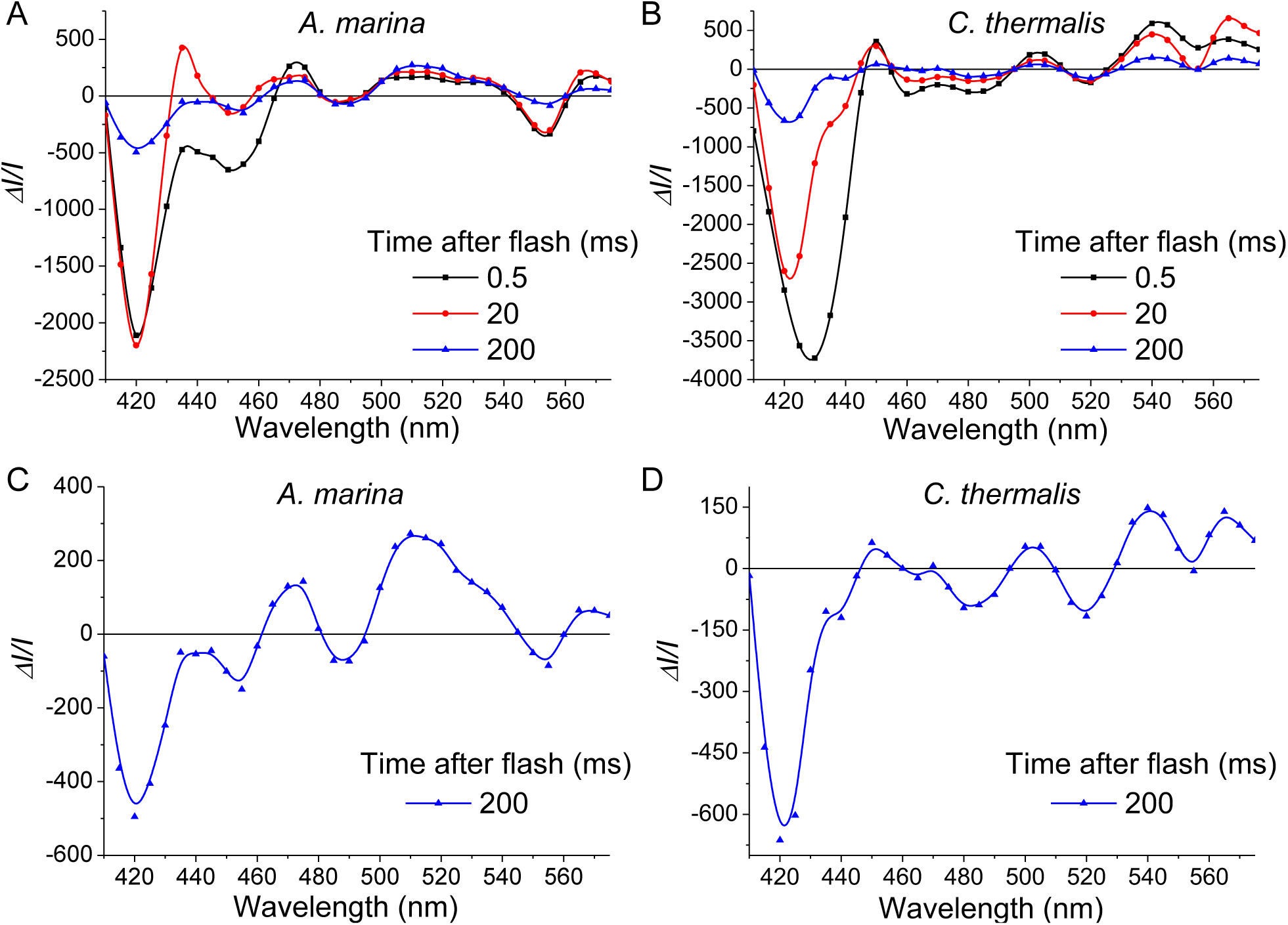
Flash-induced difference absorption spectra of *A. marina* and *C. thermalis* in the absence of PMS. (A) and (B) Spectra of flash-induced absorption changes in *A. marina* and *C. thermalis* cells, respectively, determined in the presence of DCMU and HA in anoxic conditions. Absorption changes were sampled at the indicated time intervals (in ms) after the flash. The spectra recorded at 200 ms after the flash are shown with an expanded y-axis scale in panels (C) and (D).

In *A. marina*, the negative features at 420 and 554 nm due to the oxidation of cyt *f* and the positive features at 435 and 563 nm due to the reduction of the *b* hemes of cyt *b*_6_*f* are present at both 500 µs and 20 ms after the flash (Fig. 4A). At 500 µs after the flash, an additional bleach at ∼450 nm and a positive feature between ∼470 and ∼500 nm can be ascribed to P_740+_. The P_A_P_B_ chlorophylls of *A. marina* PSI are both Chl *d*, and therefore have a red-shifted absorption in the Soret with respect to its Chl *a* counterpart[49]. At 500 µs and 20 ms after the flash, the *C. thermalis* spectra also contain features corresponding to the oxidation of cyt *f* and reduction of the *b* hemes of cyt *b*_6_*f*, although the latter are less well resolved in the Soret as they superimpose on absorption changes from P_700_^+^. The contribution of P_700_^+^ dominates in the Soret at 500 µs after the flash, with a negative peak at 430 nm and a positive feature at ∼445 nm.

Like in *Synechocystis* and *S. elongatus*[41], the spectrum of P_700_^+^ does not interfere with the carotenoid ECS in the 480-500 nm region in *C. thermalis*, confirming that in this species ECS can be measured as *ΔI/I* 500 - 480 nm. In *A. marina*, instead, the positive shoulder of P_740+_ in the Soret has a higher absorption towards 480 than 500 nm, so it cannot be subtracted from ECS using this combination of wavelengths. On the other hand, since the absorption spectrum of cyt *f* is not different in *A. marina*, ECS can be measured in this species as *ΔI/I* 520-546 nm, as in plants and green algae.

Having determined the appropriate combination of wavelengths for their measurement, we tested whether the ECS signals of the two far-red cyanobacteria are linearly proportional to the trans-thylakoid Δψ, as previously demonstrated in *S. elongatus* and *Synechocystis*[41]. We measured the ECS and P_700_^+^/P_740_^+^ kinetics induced by two single-turnover saturating flashes fired at a time interval of 50 ms.

In the presence of DCMU and HA, PSII is fully inhibited and the Δψ measured after a flash corresponds to 1 charge separation per PSI. In anoxic conditions Δψ decays slowly after the flash (Fig. S2), similar to other photosynthetic organisms, because the ATP synthase slows down at the low *pmf* induced by a block of respiration in the dark[53]. In contrast, P_700+_/P_740_^+^ is re-reduced more rapidly[41], and the thus re-opened PSI generates additional Δψ on a second flash. If the ECS signals are linearly proportional to Δψ, the increase in ECS on the second flash should be equal to the first flash in proportion to the fraction of PSI being re-opened during the interval between the two flashes.

This was confirmed for *S. elongatus* (Fig. 5A and D). In *A. marina* (Fig. 5B and E) and *C. thermalis* (Fig. 5C and F), despite some variability in the ECS and P_700+_/P_740+_ decay kinetics between biological replicates, the increase in ECS induced by the second flash was also proportional to the fraction of re-opened PSI centres. Altogether, the data show a linear dependency between ECS amplitude and PSI charge separation and, therefore, transmembrane Δψ (Fig. S3).

**Fig. 5.**
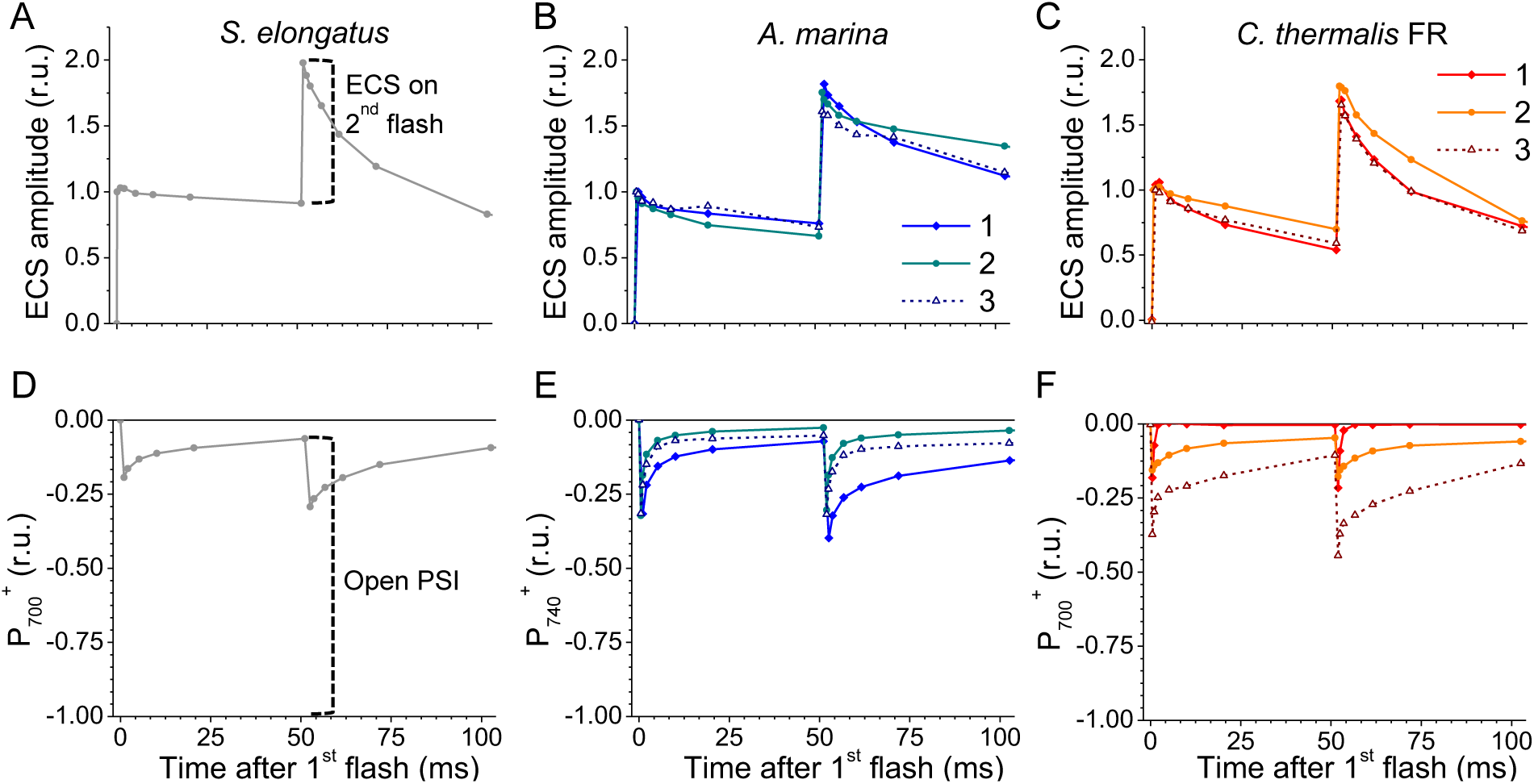
Correlation between the amplitude of flash-dependent ECS and the fraction of open PSI. Kinetics of ECS (top panels) and P_700+_/P_740_^+^ (bottom panels) signals induced by two flashes fired at a 50 ms interval measured in *S. elongatus* (A), *A. marina* (B) and *C. thermalis* (C) cells in the presence of DCMU and HA in anoxic conditions. For *A. marina* and *C. thermalis*, the results obtained in 3 biological replicates are shown. ECS signals were measured as the absorption difference between 505 and 480 nm in *S. elongatus* and *C. thermalis*, and between 520 and 546 nm in *A. marina*. All ECS traces are normalised on the amplitude at 500 µs after the first flash. P_700+_ signals were measured in D, E, and F as the absorption difference between 705 and 740 nm in *S. elongatus* and *C. thermalis*, P_740_^+^ signals at 740 nm in *A. marina*. All P_700_^+^/P_740_^+^ traces are normalised on the total amounts of P_700_^+^/P_740_^+^, measured via a multi-turnover saturating pulse in oxygenated conditions. Top and bottom panels have the same x-axis. The dashed vertical lines in panels (A) top and bottom represent, respectively, the additional ECS amplitude induced by a second flash fired 50 ms after the first and the fraction of P_700_^+^/P_740_^+^ that is re-reduced between the two flashes (i.e., the open PSI).

Of note, we observed a contribution of variable chlorophyll fluorescence (F_V_) when detecting absorption changes in the wavelength range of 705-740 nm in *C. thermalis* (Fig. S4 and S5, and associated text). This contribution was cancelled with DCMU and HA inducing maximum fluorescence F_M_ (steadily reset to baseline), so the levels of P_700_^+^ can be measured reliably as *ΔI/I* 705-740 nm in *C. thermalis* only in these conditions.

Being linearly proportional to the Δψ, the ECS signals can be used to measure the relative amounts of active PSI and PSII centres *in vivo*. The fast phase of ECS increase induced by a single-turnover saturating flash corresponds to the transmembrane Δψ generated by one charge separation per PSI in the presence of DCMU and HA, and to one charge separation per PSII+PSI in the absence of the inhibitors. Using this approach, we measured the ratio between the two photosystems in *A. marina* and *C. thermalis* (Fig. 6). The PSI/PSII ratio was on average higher in *A. marina* than in *C. thermalis* (4.3±1.6 and 1.9±0.5, respectively, in 3 biological replicates), despite the quite large variability between biological replicates

**Fig. 6.**
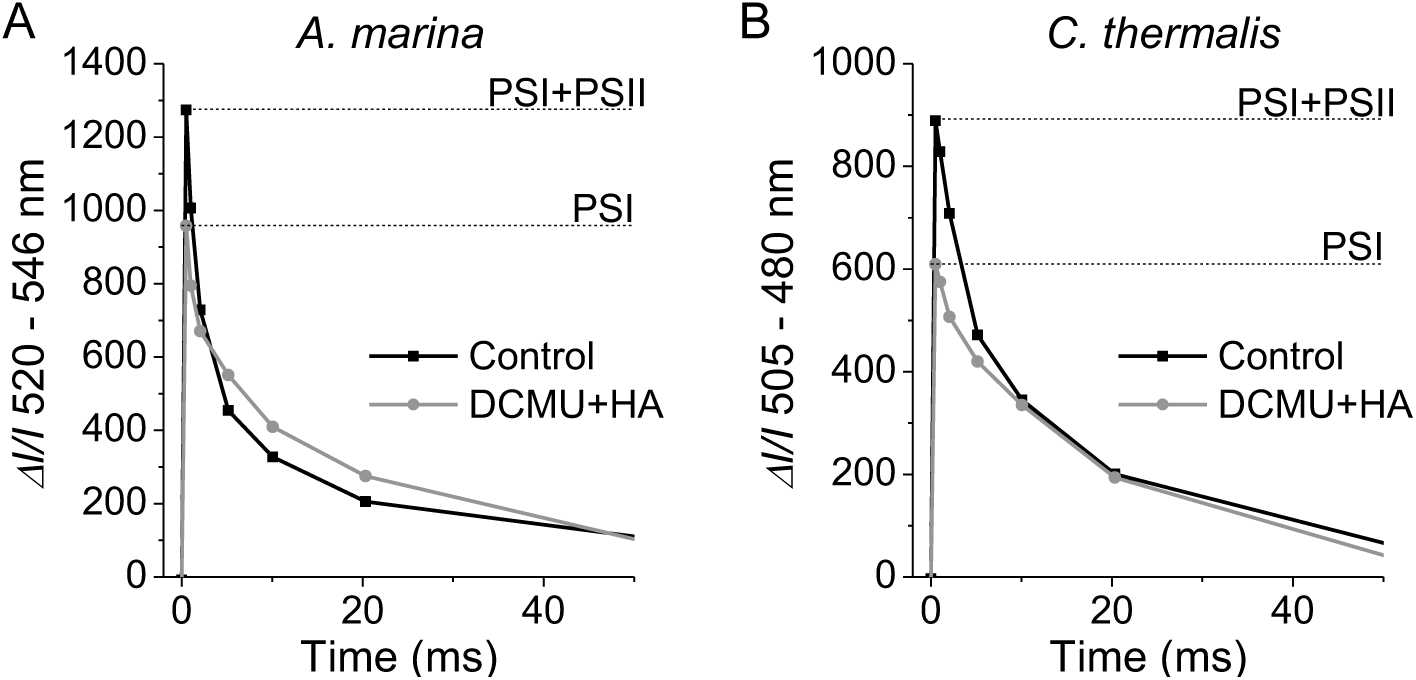
Relative amounts of active PSI and PSII measured *in vivo* by ECS. (A) and (B) Representative kinetics of flash-induced ECS signals in *A. marina* and *C. thermalis* cells, respectively, without (Control) and with the addition of PSII inhibitors (DCMU and HA).

### 3.3 ECS-based measurements of light absorption and conversion efficiency

As mentioned in the introduction, the efficiency with which light of different wavelengths induces electron transport depends on: i) the absorption cross-section (A) of the photosystems at those wavelengths, ii) the maximum efficiency with which the absorbed photons are used by the photosystems to generate a stable charge separated state (photochemical quantum yield, Φ), and iii) the operating efficiency of the electron transport chain, taking into account the limitations of the photosystems from the donor or acceptor side. The steady-state electron transport in the chain depends on several factors, including the relative amounts of PSI and PSII and the balance in excitation between the two.

ECS signals and kinetics can be used to estimate the light conversion efficiency *in vivo*, quantifying the electron flow rates and the amount of photosystems. The maximal electron transport rate, called *I* (in e^-^ PS^-1^ s^-1^) is measured as the initial rate of ECS increase at the onset of illumination, when the photosystems are open, *i.e.* charge separation is not limited by the availability of electron acceptors (for both PSII and PSI) or donors (for PSI). For a given light irradiance (Irr), *I* = Irr * A * Φ. Therefore, although ECS cannot measure individually the absorption cross-section (A) and the photochemical yield (Φ) of the photosystems, it can measure the combination of the two, *i.e.* their functional antenna size. The steady-state electron transport rate under continuous illumination, called *J* (also in e^-^ PS^-1^ s^-1^) is limited by electron transport steps downstream of the primary photochemistry in PSII and PSI. When both photosystems are active, *J* measures the combined rates of Linear and Cyclic Electron Flow (LEF and CEF, respectively).

Mathiot and Alric (2021)[17] introduced the *J*/*I* ratio as the “transmission coefficient” (a dimensionless parameter), meaning the fraction of primary photochemical events that contributes to the flow of electrons along the whole chain. The transmission coefficient can theoretically approach 1 at very low light intensities, when almost all photosystems remain open in the light. It then progressively decreases at higher light intensities as the flow of electrons extracted from the primary donors (*I*) exceeds the transport capacity of the chain (*J*). Here, we measured these ECS parameters in *S. elongatus*, *A. marina* and *C. thermalis* cells illuminated with either red or far-red light, using two LEDs centred at 658 and 732 nm, respectively (Fig. S6).

We measured *I* as a function of irradiance, both in the absence (*I*_PSII+PSI_) and in the presence (*I*_PSI_) of PSII inhibitors. The irradiances of the two actinic LEDs was measured using a photometer equipped with a radiometric detector having a flat response in the 450 to 900 nm range. The *I* values, normalised on the total PSII+PSI amounts (single turnover flash), show a linear dependency on irradiance for both 658 and 732 nm illumination in *A. marina* and *C. thermalis* (Fig. 7C and E). This linear dependency indicates that *I* is not limited by either the capacity of the photosystems to absorb photons or their charge separation rates. The slope of the linear fitting of *I*_PSII+PSI_ as a function of irradiance (Table S1) is almost as high (∼90%) for the 732 nm illumination (*I*^732^) as for the 658 nm illumination (*I*^658^) in *A. marina*, indicating an almost equal total antenna size for far-red and red light. In contrast, the slope of *I*^732^ is lower than the slope of *I*^658^ in *C. thermalis*, indicating a ∼50% smaller antenna size for far-red than red light. The ratio between the total far-red and red antenna size (slope *I*^732^ / slope *I*^658^) is thus ∼1.9 times bigger in *A. marina* than in *C. thermalis*. For *S. elongatus* (Fig. 7A), *I*^732^ remains very low with respect to *I*^658^ at all irradiances, due to the small absorption cross-section of the Chl *a* photosystems for the far-red light used here (∼0.17% of that for the red light, Table S1). The difference in the ratios between red and far-red antenna sizes between the three strains are consistent with the absorption spectra of the cells (Fig. S6).

**Fig. 7.**
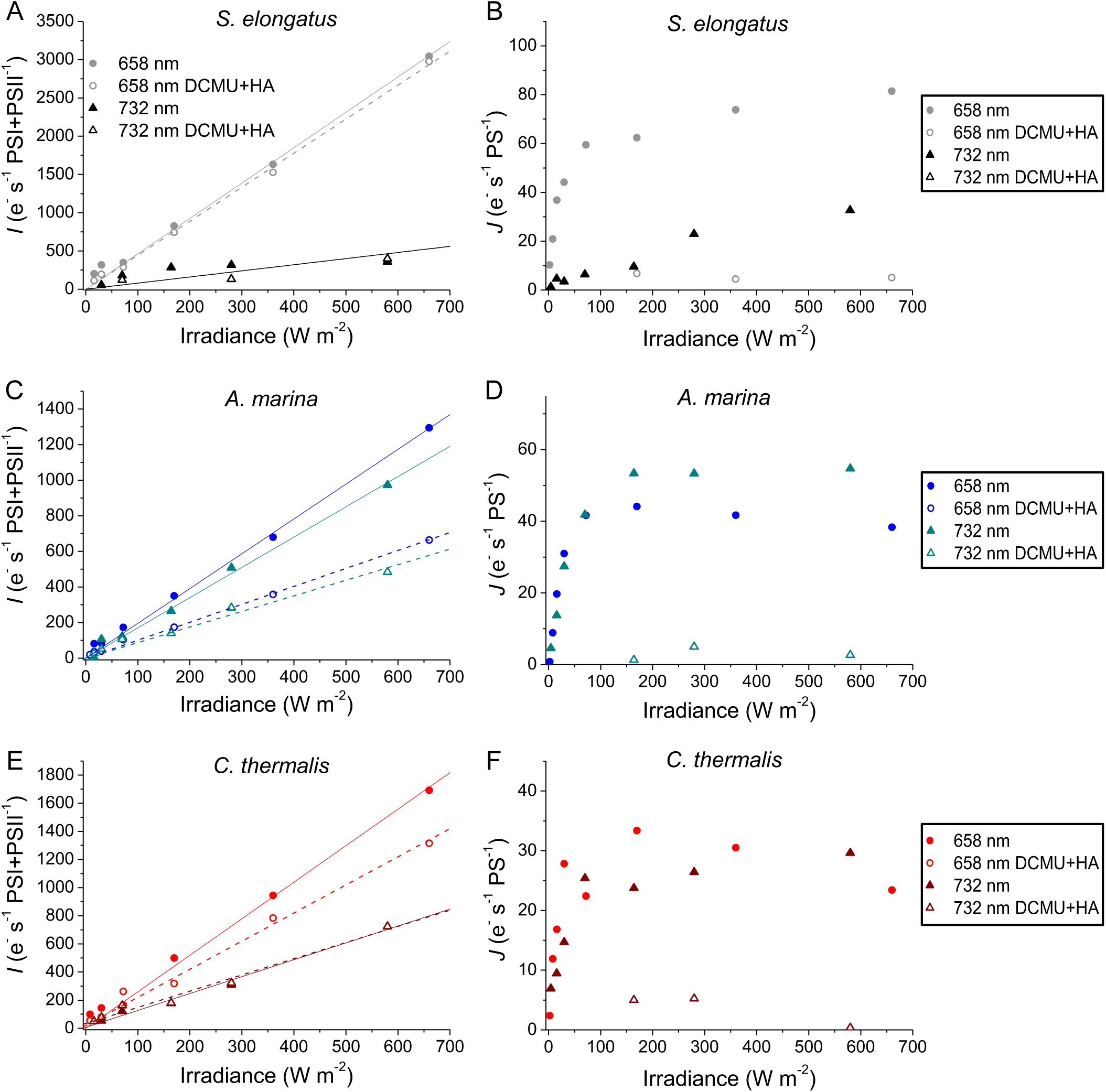
ECS-based measurements of electron transport rates as a function of light irradiance and wavelength. Maximal (*I*, panels A, C and E) and steady-state (*J*, panels B, D and F) electron transport rates measured in *S. elongatus* (A and B), *A. marina* (C and D) and *C. thermalis* (E and F) cells by ECS. *I* rates were measured based on a 200 µs illumination of the indicated wavelength and irradiance both in the absence and presence of DCMU and HA. All *I* rates were normalised on the total amounts of PSI+PSII (flash-dependent ECS without PSII inhibitors). The solid and dashed lines represent the fits of the data obtained in the absence and presence of PSII inhibitors, respectively. *J* rates were measured based on a 5 ms dark relaxation of ECS during continuous illumination of the indicated wavelength and irradiance both in the absence and presence of DCMU and HA. The *J* rates were normalised on the total amounts of active photosystems present in the two conditions (PSI+PSII in control conditions, PSI only in the presence of PSII inhibitors). For all panels: closed and open circles indicate the rates measured with the 658 nm LED in the absence and presence of DCMU and HA, respectively; closed and open triangles indicate the rates measured with the 732 nm LED in the absence and presence of DCMU and HA, respectively.

Of note, the size of the far-red absorbing antenna would be overestimated when measuring the far-red light intensity using a Photosynthetically Active Radiation (PAR) detector, whose sensitivity typically decreases outside of the 400 to 700 nm visible range. Since this overestimation would be equal for all samples, though, the quantitative comparison between species of the ratio between the far-red and red antenna sizes measured by ECS would not change (i.e., slope *I*^732^ / slope *I*^658^ would remain ∼1.9 times bigger in *A. marina* than in *C. thermalis* from Fig. 7), irrespectively of how light intensities are measured.

The *I* rates also allow us to estimate the ratio between the antenna size per PSI and PSII reaction centre for the two illumination wavelengths (Table S1). For each wavelength, we normalise the slopes of *I*_PSI_ and *I*_PSII_ (the latter calculated as *I*_PSII+PSI_ – *I*_PSI_) on the relative amounts of PSI and PSII ([PSI] and [PSII]), respectively, and then calculate the ratio between the two: (slope *I*_PSI_ / [PSI]) / (slope *I*_PSII_ / [PSII]). In the case of *A. marina*, both the red (658 nm) and far-red (732 nm) antenna sizes per reaction centre are smaller for PSI than for PSII (∼0.75 each for the sample used in Fig. 7C). Since PSI cores contain ∼3 times more chlorophyll per reaction centre than PSII, these results indicate that the extrinsic antennas (both PBS-Vis and Chl *d*-containing CP43-like) are predominantly connected with PSII *in vivo*, increasing its functional antenna size for both red and far-red light.

In *C. thermalis*, the red antenna size per reaction centre is also smaller in PSI than in PSII (∼0.72 for the sample used in Fig. 7E), although Chl *f*-PSI cores also contain ∼3 times more Chl *a* molecules than Chl *f*-PSII cores[11]. This might be taken as suggesting that PBS-Vis are predominantly associated with PSII, although there is evidence that they are at least partially disconnected from the Chl *f* photosystems in *C. thermalis* cells[46]. The far-red antenna size of PSI is instead ∼4.7 times bigger than that of PSII. Since the number of far-red chlorophylls per reaction centre is only 1.6 times higher in Chl *f*-PSI than in Chl *f*-PSII (8 and 5, respectively [11]), this might suggest that PBS-FR are predominantly associated with PSI in our experimental conditions.

In *S. elongatus*, the red antenna size per reaction centre is ∼4 times bigger in PSI than in PSII for the sample used in Fig. 7A. This likely reflects that, in addition to the ∼3 times higher chlorophyll content of the cores, PBS-Vis are more associated to PSI. This is consistent with the cells being in state 2 (antennas associated to PSI) more than in state 1 (antennas associated to PSII) in the dark-adapted state under which these measurements were performed, as it is normally the case for these and other cyanobacteria[54]. In the case of the 732 nm light, the negligible difference between the *I*_PSII+PSI_ and *I*_PSI_ values indicate that it is predominantly absorbed by PSI, as expected.

ECS-based *I* rates can provide information on how excitation with different light wavelengths is distributed between the two photosystems in the different species. However, due to the low signal-to-noise of the measurements and small differences between *I*_PSII+PSI_ and *I*_PSI_ slopes in samples having a high PSI/PSII ratio, several biological replicates would be required for a robust quantitative comparison. Additionally, this method measures the antenna sizes in the dark-adapted state. Many cyanobacteria, including *S. elongatus*, are in state 2 in the dark and undergo state transitions upon illumination[54]. State transitions in *A. marina* and *C. thermalis* are yet to be investigated, but their relative PSI and PSII antenna sizes could also change upon illumination, although the total *I*_PSII+PSI_ may not.

Fig. 7B, D and F show the steady-state electron transport rates (*J*) measured under continuous illumination in the same samples as in Fig. 7A, C and E, plotted as a function of irradiance. The rates were measured in the absence of DCMU and HA (*J*_PSII+PSI_) for both 658 nm and 732 nm illumination in all three species, and in the presence of the PSII inhibitors (*J*_PSI_) with the 658 nm illumination in *S. elongatus* and with the 732 nm illumination in *A. marina* and *C. thermalis*. In all strains, *J*_PSI_ is low (∼10%) compared to *J*_PSII+PSI_, and saturates at low irradiances as already shown for *S. elongatus* and *Synechocystis*[41]. This indicates that also in *A. marina* and *C. thermalis* electron transport around PSI is limited in the absence of electron input from PSII to the intersystem chain.

When both photosystems are active, *J*_PSII+PSI_ increases similarly as a function of irradiance with both 658 nm and 732 nm illumination in *A. marina* and *C. thermalis*, saturating at ∼50 e^-^ s^-1^ PS^-1^ and ∼30 e^-^ s^-1^ PS^-1^, in the respective species.

In *S. elongatus*, *J*_PSII+PSI_ with 658 nm illumination saturates at ∼80 e^-^ s^-1^ PS^-1^, while *J*_PSII+PSI_ with 732 nm illumination progressively increases up to ∼35 e^-^ s^-1^ PS^-1^ at the highest irradiance, without reaching saturation. The maximal *J*^732^ (∼ 30 e^-^.s^-1^.PS^-1^) in Fig. 7B are well below the maximal *I*^732^ (∼ 300 e^-^.s^-1^.PS^-1^) in Fig. 7A, showing that the former are not limited by light absorption but could result from an imbalance in excitation between PSII and PSI. Indeed, the increase of *J*^732^ as a function of irradiance after the initial lag phase is comparatively faster than the increase of *I*^732^, which becomes very evident when *J* is plotted as a function of *I* (Fig. S8A). At low intensities of 732 nm illumination, PSI gets excited specifically, and *J* is limited by electron input to the chain by PSII, as seen when DCMU and HA are present. At higher irradiances, a non-negligible fraction of PSII gets excited by the shorter wavelength tail of the emission of the 732 nm LED (Fig. 6A), too small to be detectable as an increase of the *I* rates but potentially feeding enough electrons to sustain a higher PSI turnover. Measurements of *I* and *J* rates were performed using independent cultures of the three cyanobacteria, obtaining qualitatively similar results (replicates can be found in Fig. S7 and results of the fits in Table S2).

To estimate the conversion efficiency of the absorbed red and far-red light into photosynthetic electron transport, we plotted the transmission coefficient *J*/*I* as a function of *I* (Fig. 8), using the *I* and *J* rates from two biological replicates for each cyanobacterium (Fig. 7 and S7). Division by small values of *I* (in practice below 50 to 100 e^-^ s^-1^ PS^-1^, depending on the dataset) induced large scattering in *J*/*I* ratio, so these values are not reported.

**Fig. 8.**
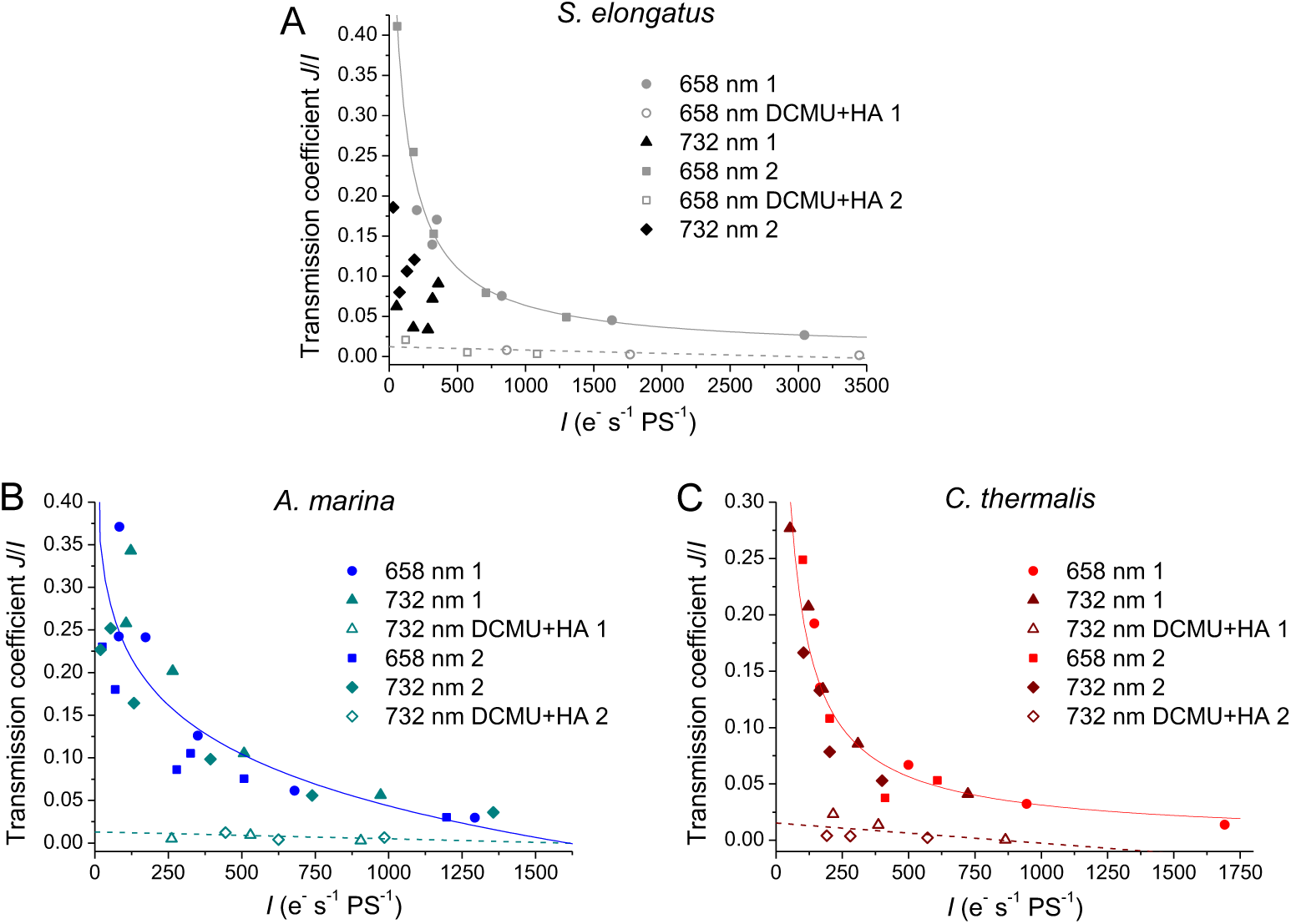
Efficiency of conversion of absorbed light into photosynthetic electron transport measured by ECS. Plots of the transmission coefficient *J*/*I* as a function of the maximal electron transport rates *I* in *S. elongatus* (A), *A. marina* (B) and *C. thermalis* (C), calculated from the data in Fig. 7 and S7 (2 biological replicates per strain). The solid lines represent the Hill fits of the *J* rates obtained in control conditions with both 658 nm and 732 nm illumination in *A. marina* and *C. thermalis*, and with 658 nm only in *S. elongatus*. The dashed lines represent the linear fits of the *J* rates obtained in the presence of DCMU and HA.

In all three species, the *J*/*I* measured in the presence of DCMU and HA remains very low (<0.025) at all *I* rates, consistent with PSI turnover being limited by the lack of electron donors in the absence of PSII activity.

In the absence of inhibitors, the *J*/*I* for the 658 nm illumination of *S. elongatus* (Fig. 8A) shows a decrease from the lowest to the highest *I* rates. This behaviour reflects the maximal rates of charge separation greatly exceeding the capacity of the rate-limiting steps of steady-state electron transport at high irradiations. The *J*/*I* for the 732 nm illumination, instead, remains always below the *J*/*I* calculated for the 658 nm illumination at corresponding *I* values, although it tends to increase at the highest *I*, after an initial decrease. This is in line with selective excitation of PSI limiting its turnover due to lack of electron donors at the lowest *I*, with the limitation being partially relieved as irradiance increases enough to excite PSII (see also Fig. S8A and B).

In *A. marina* and *C. thermalis*, the *J*/*I* for both the 658 nm and 732 nm illumination shows the same exponential decrease as a function of *I*. This indicates that no imbalance in excitation of PSI and PSII occurs under the far-red light used here, and steady-state electron transport is constrained by the same rate-limiting step(s) as in red light. This is consistent with the PSI and PSII of each species having similar absorption in the region covered by the 732 nm LED emission (Fig. 2C). Altogether, our results show that while *S. elongatus* is less efficient in converting far-red than red light into electron transport, the conversion efficiency is similar for the two wavelengths in both far-red absorbing cyanobacteria. However, the small amplitude of the ECS signals used to measure *I* and *J* rates at the lowest light irradiances prevents the comparison of the maximal conversion efficiency between strains.

## 4 Conclusions

Here we have provided the first characterisation of ECS signals in the far-red cyanobacteria *A. marina* and *C. thermalis*, also highlighting how differences in pigment composition should be taken into account when transposing absorption spectroscopy measurements between different species. We have shown that in *A. marina* and *C. thermalis*, ECS can be used to quantify the relative amounts of PSI and PSII *in vivo* as well as the relative size of their antennas. Despite some technical limitations, the ECS-based method allows for a direct and quantitative comparison of the photosynthetic light conversion efficiency as a function of illumination wavelength between species, irrespective of the absolute light intensities measured for the different wavelengths.

Our results indicate that, in contrast to Chl *a*-based cyanobacteria, both Chl *d*- and Chl *f*-containing cyanobacteria use 732 nm far-red light as efficiently as red light, despite their considerably different pigment composition. These species thus do not suffer from the excitation imbalance between PSI and PSII induced by far-red light in Chl *a*-based cyanobacteria. This result is perhaps not surprising, but it can serve as the starting point for addressing new questions, like exploring a new “far-red limit” for oxygenic photosynthesis. This limit might occur at longer wavelengths in *C. thermalis* than in *A. marina*, because of the more red-shifted Chl *f* antenna of its photosystems. However, it could be constrained by excitation imbalance between the two photosystems, since the antenna of Chl *f*-PSI extends to considerably longer wavelengths than that of Chl *f*-PSII. The excitation imbalance could be different in other Chl *f*-containing cyanobacteria whose photosystems vary in terms of number and peak absorption wavelength of the far-red chlorophylls (*e.g.* [30]). In *C. thermalis* and other species reversibly switching between Chl *a* and Chl *f* photosystems, the use of different light wavelengths could also change during the transitions between growth in visible and far-red light, *i.e.* under conditions in which the replacement of PSI, PSII, and PBS isoforms might not occur at the same rates.

In conclusion, ECS measurements on their own do not provide all the answers, but they constitute a new tool to assess the efficiency of naturally occurring far-red photosynthesis. A better understanding of the natural far-red systems, besides shedding new light on the energetic limits of oxygenic photosynthesis, could be beneficial to inform strategies aimed at expanding light usage in other organisms of interest such as crop plants or microalgae.

## Author contributions

Stefania Viola: conceptualisation, investigation, formal analysis, project administration, funding acquisition, writing – original draft. Julien Sellés: conceptualisation, investigation, resources, writing – review & editing. Jean Alric: conceptualisation, methodology, formal analysis, resources, writing – review & editing. Geoffry A. Davis: investigation, writing – review & editing. A. William Rutherford: conceptualisation, funding acquisition, writing – review & editing.

## Acknowledgments

This work received support from the French government under the France 2030 investment plan, as part of the Initiative d’Excellence d’Aix-Marseille Université - A*MIDEX and is part of the Institute of Microbiology, Bioenergies and Biotechnology - IM2B (AMX-19-IET-006). This work was also supported by the Biotechnology and Biological Sciences Research Council (BB/R001383/1, BB/V002015/1 and BB/R00921X), the Leverhulme Trust (RPG-2022-203), the Royal Society (Royal Society Research Professorship 2024), the Agence Nationale de la Recherche (CyanoFlow: ANR-23-CE20-0009) and the LabEx DYNAMO (ANR-11-LABX-0011-01).

## References

[1] X.-G. Zhu, S.P. Long, D.R. Ort, What is the maximum efficiency with which photosynthesis can convert solar energy into biomass?, Curr. Opin. Biotechnol. 19 (2008) 153–159. 10.1016/j.copbio.2008.02.004.

[2] R.E. Blankenship, Molecular mechanisms of photosynthesis, Blackwell Science, Oxford, 2008.

[3] W. Martin, K. Kowallik, Annotated English translation of Mereschkowsky’s 1905 paper ‘Über Natur und Ursprung der Chromatophoren imPflanzenreiche,’ Eur. J. Phycol. 34 (1999) 287–295. 10.1080/09670269910001736342.

[4] J.A. Raven, J.F. Allen, Genomics and chloroplast evolution: What did cyanobacteria do for plants?, Genome Biol. 4 (2003) 1–5. 10.1186/gb-2003-4-3-209.

[5] N. Nelson, C.F. Yocum, STRUCTURE AND FUNCTION OF PHOTOSYSTEMS I AND II, Annu. Rev. Plant Biol. 57 (2006) 521–565. 10.1146/annurev.arplant.57.032905.105350.

[6] R. Croce, H.V. Amerongen, Natural strategies for photosynthetic light harvesting, Nat. Chem. Biol. 10 (2014) 492–501. 10.1038/nchembio.1555.

[7] G.H. Schatz, H. Brock, A.R. Holzwarth, Kinetic and energetic model for the primary processes in photosystem II, Biophys. J. 54 (1988) 397–405. 10.1016/S0006-3495(88)82973-4.

[8] B. Gobets, R. van Grondelle, Energy transfer and trapping in photosystem I, Biochim. Biophys. Acta BBA - Bioenerg. 1507 (2001) 80–99. 10.1016/S0005-2728(01)00203-1.

[9] M. Chen, R.E. Blankenship, Expanding the solar spectrum used by photosynthesis, Trends Plant Sci. 16 (2011) 427–431. 10.1016/j.tplants.2011.03.011.

[10] A.W. Rutherford, A. Osyczka, F. Rappaport, Back-reactions, short-circuits, leaks and other energy wasteful reactions in biological electron transfer: Redox tuning to survive life in O 2, FEBS Lett. 586 (2012) 603–616. 10.1016/j.febslet.2011.12.039.

[11] D.J. Nürnberg, J. Morton, S. Santabarbara, A. Telfer, P. Joliot, L.A. Antonaru, A.V. Ruban, T. Cardona, E. Krausz, A. Boussac, A. Fantuzzi, A.W. Rutherford, Photochemistry beyond the red limit in chlorophyll f–containing photosystems, Science 360 (2018) 1210–1213. 10.1126/science.aar8313.

[12] N.V. Karapetyan, A.R. Holzwarth, M. Rögner, The photosystem I trimer of cyanobacteria: molecular organization, excitation dynamics and physiological significance, FEBS Lett. 460 (1999) 395–400. 10.1016/S0014-5793(99)01352-6.

[13] R. Croce, H. Van Amerongen, Light-harvesting in photosystem I, Photosynth. Res. 116 (2013) 153–166. 10.1007/s11120-013-9838-x.

[14] C. Wilhelm, T. Jakob, Uphill energy transfer from long-wavelength absorbing chlorophylls to PS II in Ostreobium sp. is functional in carbon assimilation, Photosynth. Res. 87 (2006) 323. 10.1007/s11120-005-9002-3.

[15] E. Kotabová, J. Jarešová, R. Kaňa, R. Sobotka, D. Bína, O. Prášil, Novel type of red-shifted chlorophyll a antenna complex from *Chromera velia*. I. Physiological relevance and functional connection to photosystems, Biochim. Biophys. Acta BBA - Bioenerg. 1837 (2014) 734–743. 10.1016/j.bbabio.2014.01.012.

[16] M. Kosugi, S.-I. Ozawa, Y. Takahashi, Y. Kamei, S. Itoh, S. Kudoh, Y. Kashino, H. Koike, Red-shifted chlorophyll a bands allow uphill energy transfer to photosystem II reaction centers in an aerial green alga, *Prasiola crispa*, harvested in Antarctica, Biochim. Biophys. Acta BBA - Bioenerg. 1861 (2020) 148139. 10.1016/j.bbabio.2019.148139.

[17] C. Mathiot, J. Alric, Standard units for ElectroChromic Shift measurements in plant biology, J. Exp. Bot. 72 (2021) 6467–6473. 10.1093/jxb/erab261.

[18] R. Emerson, C.M. Lewis, THE DEPENDENCE OF THE QUANTUM YIELD OF CHLORELLA PHOTOSYNTHESIS ON WAVE LENGTH OF LIGHT, Am. J. Bot. 30 (1943) 165–178. 10.1002/j.1537-2197.1943.tb14744.x.

[19] A. Murakami, H. Miyashita, M. Iseki, K. Adachi, M. Mimuro, Chlorophyll d in an epiphytic cyanobacterium of red algae, Science 303 (2004) 1633. 10.1126/science.1095459.

[20] H. Miyashita, H. Ikemoto, N. Kurano, K. Adachi, M. Chihara, S. Miyachi, Chlorophyll d as a major pigment, Nature 383 (1996) 402–402. 10.1038/383402a0.

[21] S. Ohashi, H. Miyashita, N. Okada, T. Iemura, T. Watanabe, M. Kobayashi, Unique photosystems in *Acaryochloris marina*, Photosynth. Res. 98 (2008) 141–149. 10.1007/s11120-008-9383-1.

[22] M. Chen, T.S. Bibby, J. Nield, A.W.D. Larkum, J. Barber, Structure of a large photosystem II supercomplex from *Acaryochloris marina*, FEBS Lett. 579 (2005) 1306–1310. 10.1016/j.febslet.2005.01.023.

[23] L. Shen, Y. Gao, K. Tang, R. Qi, L. Fu, J.-H. Chen, W. Wang, X. Ma, P. Li, M. Chen, T. Kuang, X. Zhang, J.-R. Shen, P. Wang, G. Han, Structure of a unique PSII-Pcb tetrameric megacomplex in a chlorophyll *d* –containing cyanobacterium, Sci. Adv. 10 (2024) eadk7140. 10.1126/sciadv.adk7140.

[24] M. Chen, M. Schliep, R.D. Willows, Z.-L. Cai, B.A. Neilan, H. Scheer, A red-shifted chlorophyll, Science 329 (2010) 1318–1319. 10.1126/science.1191127.

[25] F. Gan, S. Zhang, N.C. Rockwell, S.S. Martin, J.C. Lagarias, D.A. Bryant, Extensive remodeling of a cyanobacterial photosynthetic apparatus in far-red light, Science 345 (2014) 1312–1317. 10.1126/science.1256963.

[26] C.J. Gisriel, G. Shen, M.Y. Ho, V. Kurashov, D.A. Flesher, J. Wang, W.H. Armstrong, J.H. Golbeck, M.R. Gunner, D.J. Vinyard, R.J. Debus, G.W. Brudvig, D.A. Bryant, Structure of a monomeric photosystem II core complex from a cyanobacterium acclimated to far-red light reveals the functions of chlorophylls d and f, J. Biol. Chem. 298 (2022) 101424. 10.1016/j.jbc.2021.101424.

[27] M. Judd, J. Morton, D. Nürnberg, A. Fantuzzi, A.W. Rutherford, R. Purchase, N. Cox, E. Krausz, The primary donor of far-red photosystem II: ChlD1 or PD2?, Biochim. Biophys. Acta BBA - Bioenerg. 1861 (2020) 148248. 10.1016/j.bbabio.2020.148248.

[28] D.A. Cherepanov, I.V. Shelaev, F.E. Gostev, A.V. Aybush, M.D. Mamedov, G. Shen, V.A. Nadtochenko, D.A. Bryant, A.Y. Semenov, J.H. Golbeck, Evidence that chlorophyll f functions solely as an antenna pigment in far-red-light photosystem I from *Fischerella thermalis* PCC 7521, Biochim. Biophys. Acta - Bioenerg. 1861 (2020) 148184. 10.1016/j.bbabio.2020.148184.

[29] C. Gisriel, G. Shen, V. Kurashov, M.Y. Ho, S. Zhang, D. Williams, J.H. Golbeck, P. Fromme, D.A. Bryant, The structure of Photosystem I acclimated to far-red light illuminates an ecologically important acclimation process in photosynthesis, Sci. Adv. 6 (2020) 1–12. 10.1126/sciadv.aay6415.

[30] M. Tros, V. Mascoli, G. Shen, M.Y. Ho, L. Bersanini, C.J. Gisriel, D.A. Bryant, R. Croce, Breaking the red limit: efficient trapping of long-wavelength excitations in chlorophyll-f-containing photosystem I, Chem 7 (2021) 155–173. 10.1016/j.chempr.2020.10.024.

[31] L.A. Antonaru, T. Cardona, A.W.D. Larkum, D.J. Nürnberg, Global distribution of a chlorophyll f cyanobacterial marker, ISME J. 14 (2020) 2275–2287. 10.1038/s41396-020-0670-y.

[32] R.E. Blankenship, D.M. Tiede, J. Barber, G.W. Brudvig, G. Fleming, M. Ghirardi, M.R. Gunner, W. Junge, D.M. Kramer, A. Melis, T.A. Moore, C.C. Moser, D.G. Nocera, A.J. Nozik, D.R. Ort, W.W. Parson, R.C. Prince, R.T. Sayre, Comparing photosynthetic and photovoltaic efficiencies and recognizing the potential for improvement, Science 332 (2011) 805–809. 10.1126/science.1200165.

[33] C.A.R. Cotton, J.S. Douglass, S.D. Causmaecker, K. Brinkert, T. Cardona, A. Fantuzzi, A.W. Rutherford, J.W. Murray, Photosynthetic constraints on fuel from microbes, Front. Bioeng. Biotechnol. 3 (2015) 1–5. 10.3389/fbioe.2015.00036.

[34] S. Viola, W. Roseby, S. Santabarbara, D. Nürnberg, R. Assunção, H. Dau, J. Sellés, A. Boussac, A. Fantuzzi, A.W. Rutherford, Impact of energy limitations on function and resilience in long-wavelength Photosystem II, eLife 11 (2022) e79890. 10.7554/eLife.79890.

[35] V. Mascoli, L. Bersanini, R. Croce, Far-red absorption and light-use efficiency trade-offs in chlorophyll f photosynthesis, Nat. Plants 6 (2020) 1044–1053. 10.1038/s41477-020-0718-z.

[36] V. Mascoli, A.F. Bhatti, L. Bersanini, H. Van Amerongen, R. Croce, The antenna of far-red absorbing cyanobacteria increases both absorption and quantum efficiency of Photosystem II, Nat. Commun. 13 (2022) 3562. 10.1038/s41467-022-31099-5.

[37] B. Gobets, I.H.M.V. Stokkum, M. Rögner, J. Kruip, E. Schlodder, N.V. Karapetyan, J.P. Dekker, R.V. Grondelle, Time-resolved fluorescence emission measurements of photosystem I particles of various cyanobacteria: A unified compartmental model, Biophys. J. 81 (2001) 407–424. 10.1016/S0006-3495(01)75709-8.

[38] E. Elias, T.J. Oliver, R. Croce, Oxygenic photosynthesis in far-red light: strategies and mechanisms, Annu. Rev. Phys. Chem. (2024). 10.1146/annurev-physchem-090722-125847.

[39] A. Murakami, S.J. Kim, Y. Fujita, Changes in photosystem stoichiometry in Response to environmental conditions for cell growth observed with the cyanophyte *Synechocystis* PCC 6714, Plant Cell Physiol. 38 (1997) 392–397. 10.1093/oxfordjournals.pcp.a029181.

[40] B. Bailleul, P. Cardol, C. Breyton, G. Finazzi, Electrochromism: A useful probe to study algal photosynthesis, Photosynth. Res. 106 (2010) 179–189. 10.1007/s11120-010-9579-z.

[41] S. Viola, B. Bailleul, J. Yu, P. Nixon, J. Sellés, P. Joliot, F.-A. Wollman, Probing the electric field across thylakoid membranes in cyanobacteria, Proc. Natl. Acad. Sci. 116 (2019) 21900– 21906. 10.1073/pnas.1913099116.

[42] R.Y. Stanier, J. Deruelles, R. Rippka, M. Herdman, J.B. Waterbury, Generic assignments, strain histories and properties of pure cultures of cyanobacteria, Microbiology 111 (1979) 1–61. 10.1099/00221287-111-1-1.

[43] B. Bailleul, X. Johnson, G. Finazzi, J. Barber, F. Rappaport, A. Telfer, The thermodynamics and kinetics of electron transfer between cytochrome *b*_6_*f* and photosystem I in the chlorophyll d-dominated cyanobacterium, *Acaryochloris marina*, J. Biol. Chem. 283 (2008) 25218–25226. 10.1074/jbc.M803047200.

[44] T. Hamaguchi, K. Kawakami, K. Shinzawa-Itoh, N. Inoue-Kashino, S. Itoh, K. Ifuku, E. Yamashita, K. Maeda, K. Yonekura, Y. Kashino, Structure of the far-red light utilizing photosystem I of *Acaryochloris marina*, Nat. Commun. 12 (2021) 2333. 10.1038/s41467-021-22502-8.

[45] C. Xu, Q. Zhu, J. Chen, L. Shen, X. Yi, Z. Huang, W. Wang, M. Chen, T. Kuang, J. Shen, X. Zhang, G. Han, A unique photosystem I reaction center from a chlorophyll *d* -containing cyanobacterium *Acaryochloris marina*, J. Integr. Plant Biol. 63 (2021) 1740–1752. 10.1111/jipb.13113.

[46] C. MacGregor-Chatwin, D.J. Nürnberg, P.J. Jackson, C. Vasilev, A. Hitchcock, M.-Y. Ho, G. Shen, C.J. Gisriel, W.H.J. Wood, M. Mahbub, V.M. Selinger, M.P. Johnson, M.J. Dickman, A.W. Rutherford, D.A. Bryant, C.N. Hunter, Changes in supramolecular organization of cyanobacterial thylakoid membrane complexes in response to far-red light photoacclimation, Sci. Adv. 8 (2022) eabj4437. 10.1126/sciadv.abj4437.

[47] E.G. Andrizhiyevskaya, T.M.E. Schwabe, M. Germano, S. D’Haene, J. Kruip, R. van Grondelle, J.P. Dekker, Spectroscopic properties of PSI–IsiA supercomplexes from the cyanobacterium *Synechococcus* PCC 7942, Biochim. Biophys. Acta BBA - Bioenerg. 1556 (2002) 265–272. 10.1016/S0005-2728(02)00371-7.

[48] T. Hiyama, B. Ke, Difference spectra and extinction coefficients of P700, BBA - Bioenerg. 267 (1972) 160–171. 10.1016/0005-2728(72)90147-8.

[49] Q. Hu, H. Miyashita, I. Iwasaki, N. Kurano, S. Miyachi, M. Iwaki, S. Itoh, A photosystem I reaction center driven by chlorophyll d in oxygenic photosynthesis, Proc. Natl. Acad. Sci. U.S. A. 95 (1998) 13319–13323. 10.1073/pnas.95.22.13319.

[50] T. Tomo, Y. Kato, T. Suzuki, S. Akimoto, T. Okubo, T. Noguchi, K. Hasegawa, T. Tsuchiya, K. Tanaka, M. Fukuya, N. Dohmae, T. Watanabe, M. Mimuro, Characterization of highly purified photosystem I complexes from the chlorophyll d-dominated cyanobacterium *Acaryochloris marina* MBIC 11017, J. Biol. Chem. 283 (2008) 18198–18209. 10.1074/jbc.M801805200.

[51] H.T. Witt, Energy conversion in the functional membrane of photosynthesis. Analysis by light pulse and electric pulse methods, Biochim. Biophys. Acta BBA - Rev. Bioenerg. 505 (1979) 355–427. 10.1016/0304-4173(79)90008-9.

[52] P. Joliot, A. Joliot, Electrogenic events associated with electron and proton transfers within the cytochrome *b*_6_/*f* complex, Biochim. Biophys. Acta BBA - Bioenerg. 1503 (2001) 369–376. 10.1016/S0005-2728(00)00232-2.

[53] P. Joliot, A. Joliot, Characterization of linear and quadratic electrochromic probes in *Chlorella sorokiniana* and *Chlamydomonas reinhardtii*, Biochim. Biophys. Acta BBA - Bioenerg. 975 (1989) 355–360. 10.1016/S0005-2728(89)80343-3.

[54] P.I. Calzadilla, D. Kirilovsky, Revisiting cyanobacterial state transitions, Photochem. Photobiol. Sci. 19 (2020) 585–603. 10.1039/c9pp00451c.

