## Supplementary figures and text for "*In vivo* ElectroChromic Shift measurements of photosynthetic activity in far-red absorbing cyanobacteria"

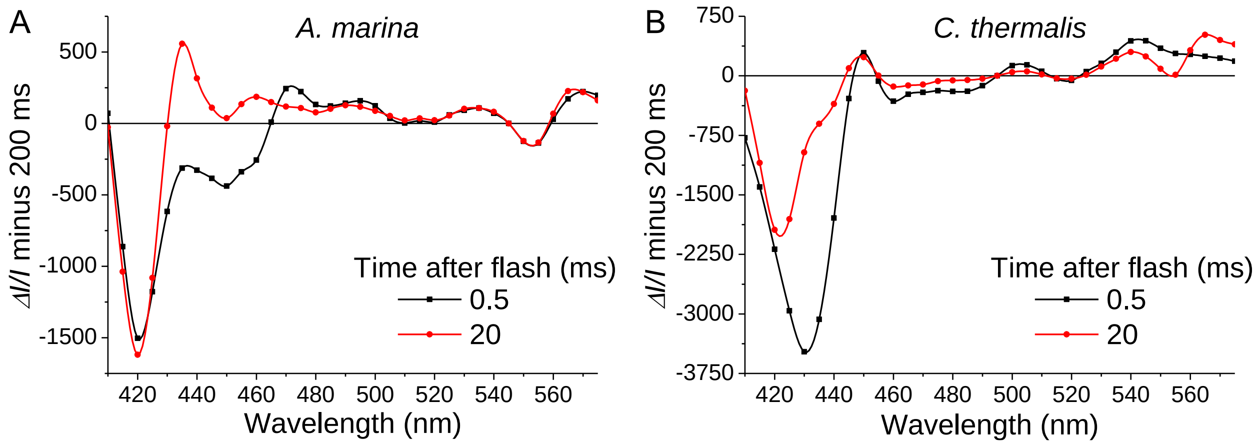


Fig. S1. Redox-dependent absorption difference signals in *A. marina* and *C. thermalis* measured in the absence of PMS. (A) and (B) Difference between the spectra recorded at 0.5 and 20 ms after the flash and those recorded at 200 ms after the flash in *A. marina* and *C. thermalis*, respectively. The differences were calculated using the spectra in Fig. 4, recorded in the presence of DCMU and HA in anoxic conditions.

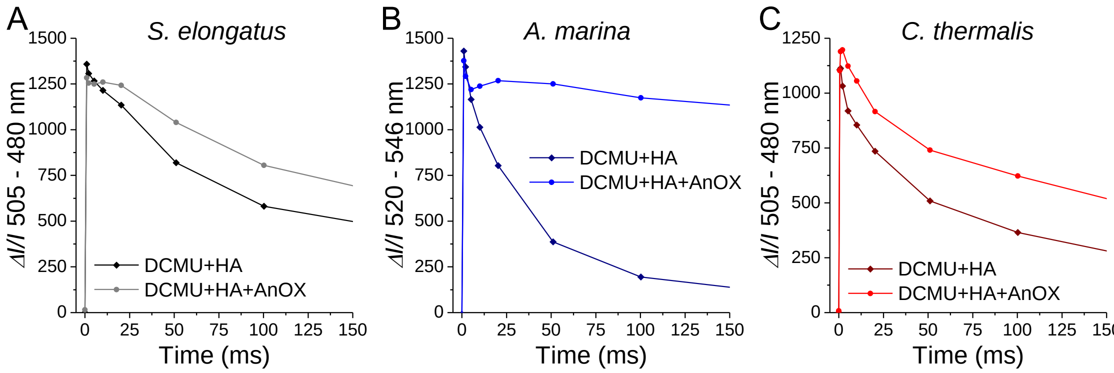


Fig. S2. Flash-induced ECS kinetics in oxygenated and anoxic conditions. Representative kinetics of flash-induced ECS signals in *S. elongatus* (A), *A. marina* (B) and *C. thermalis* (C) cells in the presence of DCMU and HA. The kinetics were recorded either in oxygenated conditions or after establishing anoxia in the measurement cuvette using glucose and glucose oxidase (AnOX). ECS signals were measured as the absorption difference between 505 and 480 nm in *S. elongatus* and *C. thermalis* and between 520 and 546 nm in *A. marina*. Only the first 150 ms of the decay kinetics are displayed here, on an expanded scale.

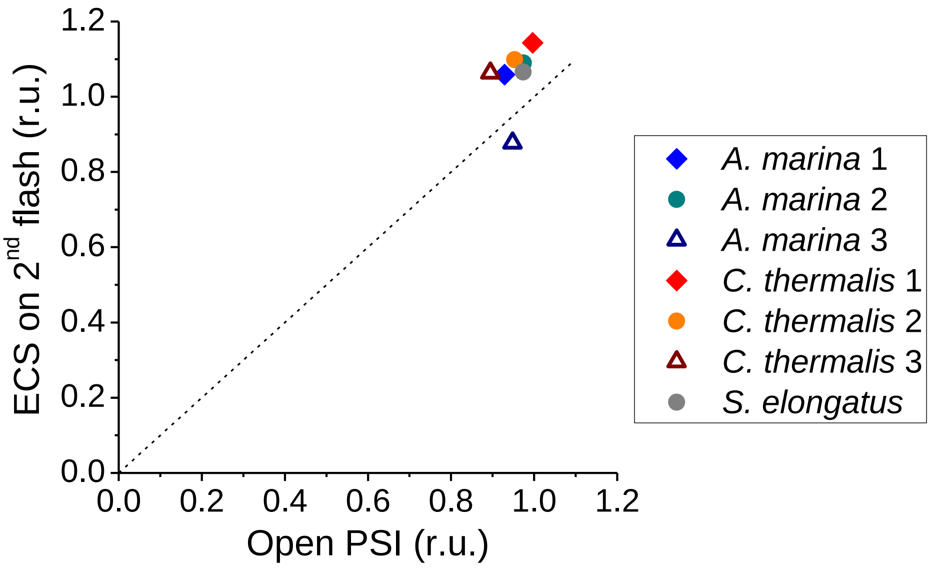


Fig. S3. Linearity of ECS signals in *A. marina* and *C. thermalis*. The additional ECS amplitude induced by a second flash fired 50 ms after the first, normalised on the amplitude of ECS induced by the first flash, is plotted against the fraction of $P_{700}^{+}/P_{740}^{+}$ that was re-reduced between the two flashes (open PSI). The values are calculated from the data in Fig. 5 (3 biological replicates each for *A. marina* and *C. thermalis*, 1 replicate for *S. elongatus*). The diagonal dashed line represents the linear dependency between x and y values.

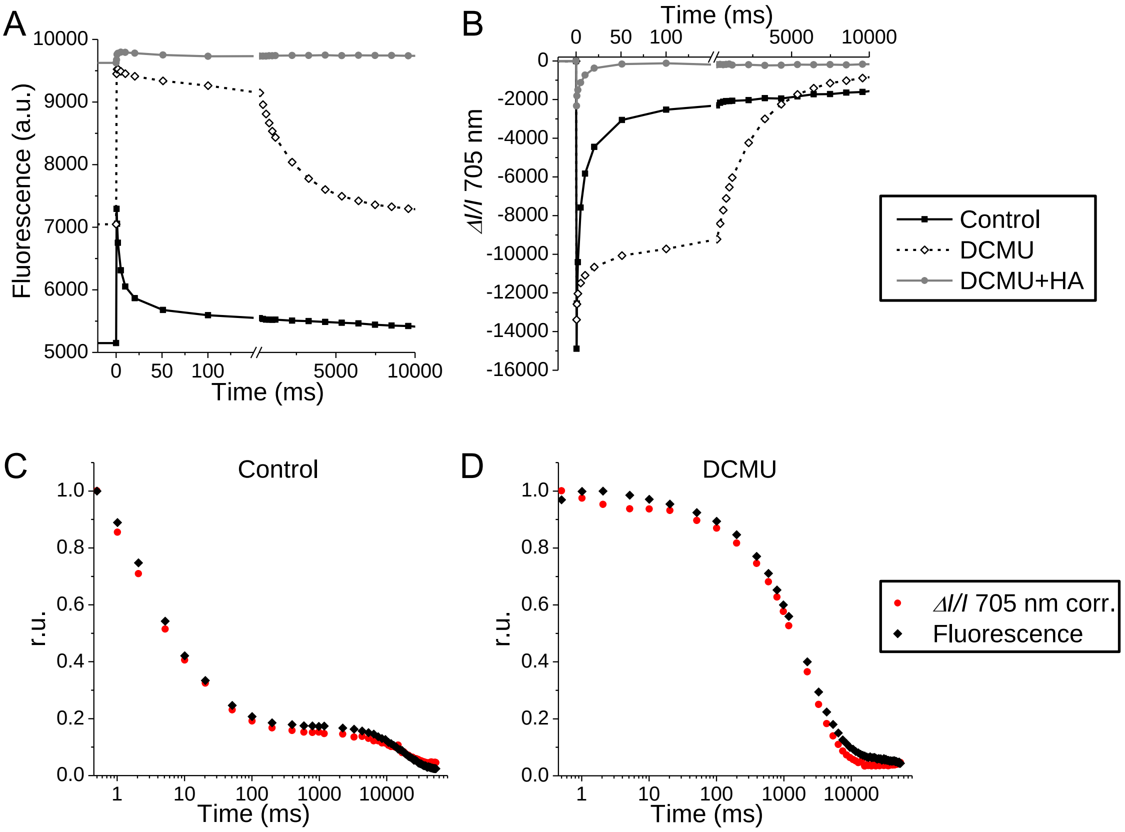


Fig. S4. The contribution of fluorescence to the absorption difference signals measured at 705 nm in *C. thermalis* cells. Decay kinetics of fluorescence (A) and of *ΔI/I* 705 nm (B) were measured after a single-turnover flash in the same *C. thermalis* cell suspension in control conditions, in the presence of DCMU only, and in the presence of both DCMU and HA. In the presence of DCMU and HA, measurements were performed after a few seconds of saturating pre-illumination that closed all PSII centres. Note that in (A) and (B) the x-axis is broken to show the first 150 ms of the decay on an expanded scale. In (C) and (D) the *ΔI/I* 705 nm decay kinetics from (B) measured in the presence of DCMU and HA were subtracted from those measured in control conditions and in the presence of DCMU only. The thus corrected *ΔI/I* 705 nm kinetics in control and DCMU are compared with the fluorescence kinetics recorded in the same conditions. Note that here the x-axis here is displayed on a log10 scale.

As stated in section 3.2 of the main text, while performing the measurements to verify the linearity of ECS, we noticed that $P_{700}^{+}$ amplitude could not be measured reliably as the difference in absorption changes at 705 and 740 nm in *C. thermalis* except when DCMU and HA were both present. Indeed, the *ΔI/I* 705 nm signals measured after a flash in the presence of

DCMU only, or in the absence of PSII inhibitors, displayed amplitudes and kinetics not compatible with those of $P_{700}^+$. Based on these observations, we hypothesized that in *C. thermalis* cells adapted to far-red light, the absorption changes measured at 705 nm contain contributions from signals linked to PSII photochemistry.

We thus compared, in the same *C. thermalis* cells, the decay kinetics of fluorescence and of *ΔI/I* 705 nm after a flash, either in the absence or in the presence of PSII inhibitors (Fig. S4A and B). As expected, the fluorescence decays with bi-phasic kinetics in control conditions, with most of the decay occurring in the μs to tens of ms time-scale due to $Q_A^-$ reoxidation *via* forward electron transfer. A slower decaying phase of smaller amplitude reflects a fraction of PSII where $Q_A^-$ reoxidation occurs *via* charge recombination[1]. In the presence of the $Q_B$ site inhibitor DCMU, $F_0$ and $F_M$ increase and the variable fluorescence decays for the most part monophasically in the seconds time-scale due to $Q_A^-$ reoxidation via charge recombination. In the presence of both PSII inhibitors, instead, $Q_A^-$ is trapped by electron donation from HA and thus cannot decay via recombination. In these conditions, variable fluorescence is suppressed because $F_0$ increases to $F_M$ levels, and PSII cannot do charge separation (*i.e.*, PSII is "closed"). The fluorescence kinetics in Fig. S4A show this behaviour.

Surprisingly, we observed a similar behaviour also for the *ΔI/I* 705 nm kinetics (Fig. S4B), with a slowly decaying phase (seconds time-scale) being already present in control conditions and predominating in the presence of DCMU. Some slowing down of $P_{700}^+$ re-reduction might be expected in the absence of electron donation by PSII, but the DCMU-dependent increase in amplitude of the slow *ΔI/I* 705 nm decay is not compatible with the kinetics of $P_{700}^+$ re-reduction after a flash previously measured in *S. elongatus* and *Synechocystis* cells[2,3]. Additionally, the presence of HA greatly reduced the amplitude of *ΔI/I* 705 nm, suppressing most of its slow decay. Since HA is not known to be a PSI electron donor, the slowly decaying component of the *ΔI/I* 705 nm measured in the absence of HA cannot be attributed to $P_{700}^+$ but instead to a component of PSII. When PSII photochemistry is inhibited by the presence of both DCMU and HA, the *ΔI/I* 705 nm should represent *bona fide* $P_{700}^+$ signals. In the far-red-adapted *C. thermalis* cells, an actinic flash at 700 nm, as used here, should primarily excite the PBS-FR, which have a broad fluorescence emission peaking around 715 nm[4]. This fluorescence could interfere with the $P_{700}^+$ measurements at 705 nm, with emission at this wavelength being detected as an apparent decrease in absorption. Indeed, the *ΔI/I* 705 nm kinetics measured in the control and in the DCMU treated sample, once corrected by subtracting the $P_{700}^+$ kinetics remaining in the presence of both DCMU and HA, matched almost exactly the fluorescence kinetics measured in the same conditions (Fig. S4C and D).

To confirm our hypothesis of fluorescence interfering with the *ΔI/I* 705 nm-based $P_{700}^{+}$ measurements in *C. thermalis*, we recorded flash-dependent absorption difference spectra in the chlorophyll Qy region in *S. elongatus*, *A. marina* and *C. thermalis* in the presence of DCMU (Fig. S5A, B and C). The spectra were recorded at 200 μs after the flash and then in the tens to hundreds of ms time-range. From our previous measurements in *S. elongatus* and *Synechocystis*[2,3], we expected ~30-40% of $P_{700}^{+}$ to remain at 200 μs after the flash, and to be fully re-reduced in the ms time-scale. In contrast, in the presence of DCMU the fluorescence is expected to follow the kinetics of $Q_A^-$ reoxidation *via* charge recombination, *i.e.* to remain stable in the ms range but to decay in hundreds of ms to seconds.

The spectra of the absorption changes decaying between 200 μs and 21 ms after the flash indeed correspond to those of $P_{700}^{+}$ in *S. elongatus* and *C. thermalis* and of $P_{740}^{+}$ in *A. marina* (Fig. S5D). As previously reported[5], the $P_{700}^{+}$ spectrum of Chl *f*-PSI of *C. thermalis* is very similar to that of Chl *a*-PSI (in this case that of *S. elongatus*), having a main bleach around 703 nm. Two additional negative features that are present in the $P_{700}^{+}$ spectrum in Chl *f*-PSI between 720 and 750 nm have been attributed to bandshifts of far-red chlorophylls located in the reaction centre or in its proximity[5,6]. On the other hand, the slowly decaying spectral component of *C. thermalis* presents two main negative peaks around 675 and 710 nm (Fig. S5C). These bleaches match the peaks of PBS-Vis and PBS-FR fluorescence emission previously recorded in *C. thermalis* cells with excitation in the green region[7] (as in our spectra, where excitation was provided by a xenon flash lamp filtered with a blue-green short-pass filter). These data show that, unless variable fluorescence is eliminated by inhibiting PSII charge separation, absorption changes at 705 nm cannot be used to measure $P_{700}^{+}$ in far-red-grown *C. thermalis*, since they are dominated by interference from PBS-FR fluorescence.

In the case of *A. marina*, we also observed a broad and prominent bleach peaking around 710 nm that decayed with kinetics compatible with charge recombination in PSII (Fig. S5B). We attribute this signal to Chl *d* fluorescence. Fortunately, the contribution of this fluorescence at 740 nm is minimal, and it does not interfere with $P_{740}^{+}$ measurements.

The slowly decaying spectral component was much smaller in amplitude, compared to $P_{700}^{+}$, in *S. elongatus* than in the two far-red species, and seemed to be mostly due to PBS-Vis fluorescence causing a negative drift towards 650 nm (Fig. S5A).

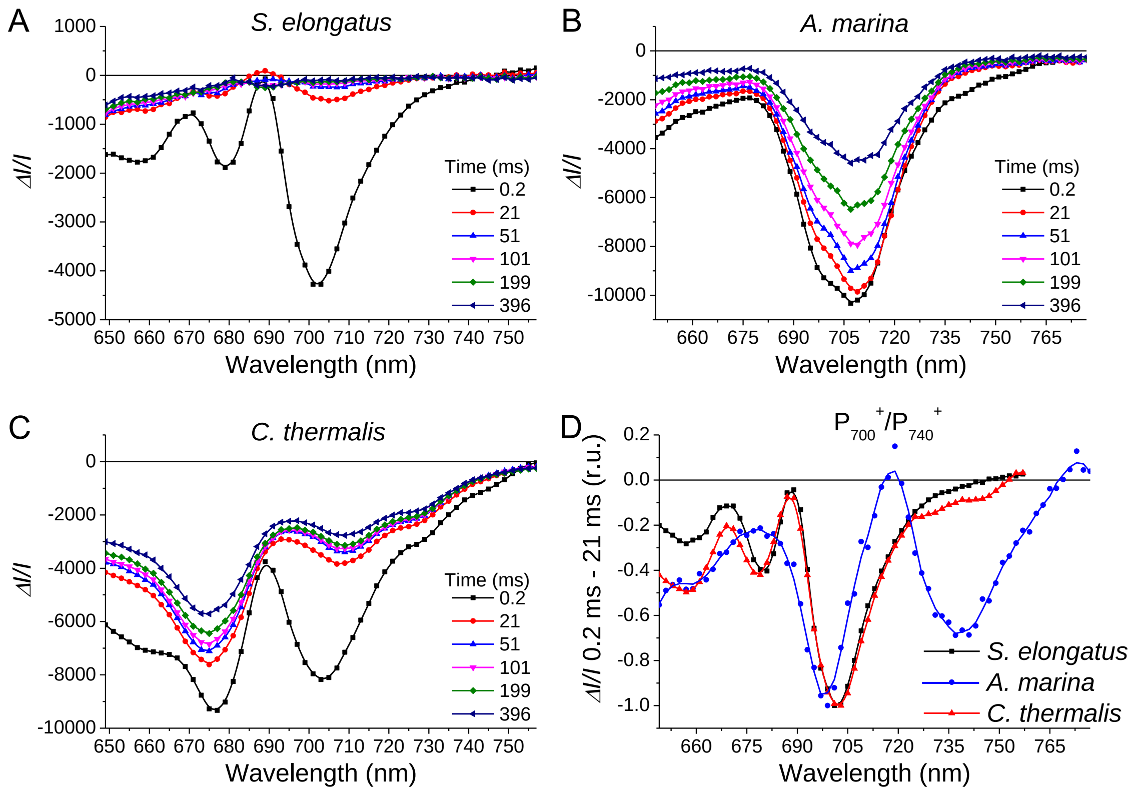


Fig. S5. Absorption difference signals in the red and far-red region measured in intact cells. (A), (B) and (C) Spectra of flash-induced absorption changes in *S. elongatus*, *A. marina* and *C. thermalis* cells, respectively, recorded in the presence of DCMU. Absorption changes were sampled at the indicated time intervals (in ms) after the flash. Note that in *A. marina* the recordings were extended 20 nm to the red with respect to *S. elongatus* and *C. thermalis*. (D) Spectra of $P_{700}^{+}$ in *S. elongatus* and *C. thermalis* and of $P_{740}^{+}$ in *A. marina*, obtained as the difference between the absorption changes measured at 0.2 and at 21 ms after the flash from panels A-C. For comparison, the $P_{700}^{+}/P_{740}^{+}$ spectra were normalised to their absorption minimum.

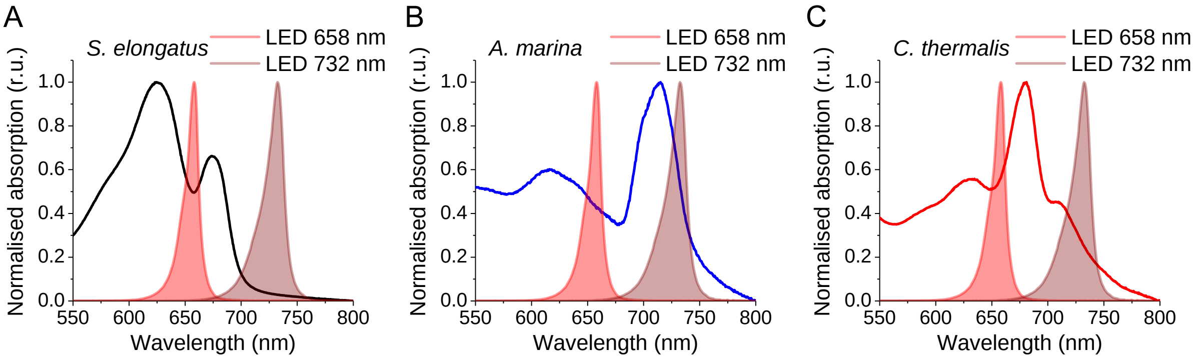


Fig. S6. Comparison between whole cell absorption spectra of the cyanobacterial species studied and the emission spectra of the actinic lights used to measure electron transport rates. The absorption spectra of *S. elongatus* (A), *A. marina* (B) and *C. thermalis* (C) cells are the same as in Fig. 1. The absorption spectra are superimposed on the emission spectra of the actinic LEDs centred at 658 nm and 732 nm used for all the measurements in section 3.3 of the main text. All spectra are normalised to their maximum in the 550 nm to 800 nm region.

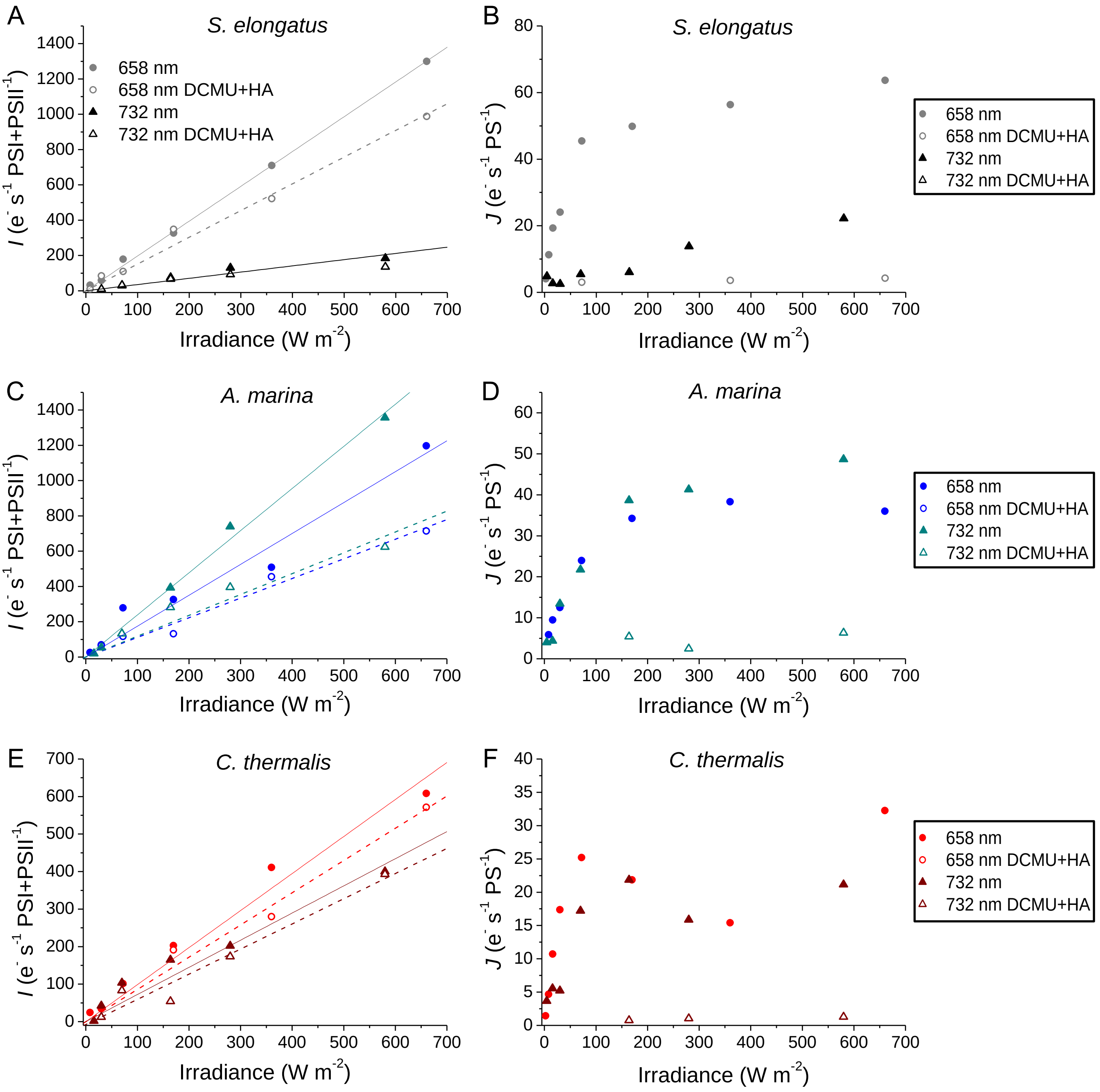


Fig. S7. ECS-based measurements of electron transport rates as a function of light irradiance and wavelength. Measurements were done as in Fig. 7, on independent biological replicates. Maximal (*I*, panels A, C and E) and steady-state (*J*, panels B, D and F) electron transport rates measured in *S. elongatus* (A and B), *A. marina* (C and D) and *C. thermalis* (E and F) cells by ECS. *I* rates were measured based on a 200 μs illumination of the indicated wavelength and irradiance both in the absence and presence of DCMU and HA. All *I* rates were normalised on the total amounts of PSI+PSII (flash-dependent ECS without PSII inhibitors). The solid and dashed lines represent the fits of the data obtained in the absence and presence of PSII inhibitors, respectively. *J* rates were measured based on a 5 ms dark relaxation of ECS during continuous illumination of the indicated wavelength and irradiance both in the absence and presence of DCMU and HA. The *J* rates were normalised on the total amounts of active photosystems present in the two conditions (PSI+PSII in control conditions, PSI only in the presence of PSII inhibitors). For all panels: closed and open circles indicate the rates measured with the 658 nm LED in the absence and presence of DCMU and HA, respectively; closed and open triangles indicate the rates measured with the 732 nm LED in the absence and presence of DCMU and HA, respectively.

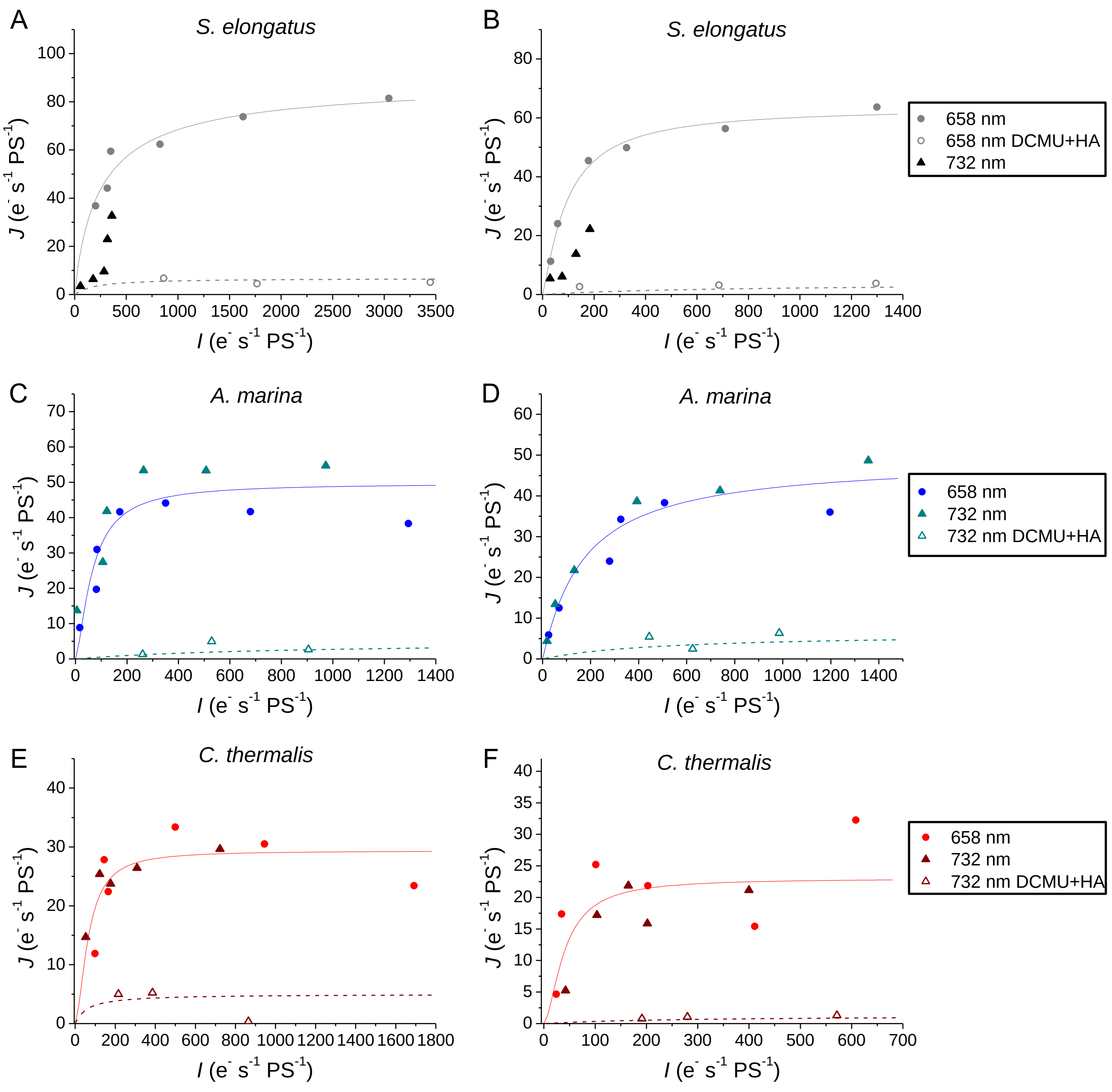


Fig. S8. ECS-based measurements of steady-state electron transport rates (*J*) as a function of maximal electron transport rates (*I*) in two biological replicates of *S. elongatus* (A and B), *A. marina* (C and D) and *C. thermalis* (E and F). The *I* and *J* rates plotted here are those from Fig. 7 (A, C and E) and Fig. S7 (B, D and F). The solid lines represent the fits of the *J* rates obtained in control conditions with both 658 nm and 732 nm illumination in *A. marina* and *C. thermalis*, and with 658 nm only in *S. elongatus*. The dashed lines represent the fits of the *J* rates obtained in the presence of DCMU and HA.

| *S. elongatus* | | | | | |
|---|---|---|---|---|---|
| [PSI]/[PS Tot] | [PSII]/[PS Tot] | PSI/PSII | | Slope $I^{732}/I^{658}$ | |
| 0.86 | 0.14 | 6.32 | | 0.17 | |
| | Slope $I_{PSII+PSI}$ | Slope $I_{PSI}$ | Slope $I_{PSI}$/[PSI] | Slope $I_{PSII}$/[PSII] | $Ant_{PSI}/Ant_{PSII}$ |
| **658 nm** | 4.62 | 4.44 | 5.15 | 1.31 | 3.93 |
| **732 nm** | 0.8 | / | / | / | / |

| *A. marina* | | | | | |
|---|---|---|---|---|---|
| [PSI]/[PS Tot] | [PSII]/[PS Tot] | PSI/PSII | | Slope $I^{732}/I^{658}$ | |
| 0.59 | 0.41 | 1.42 | | 0.87 | |
| | Slope $I_{PSII+PSI}$ | Slope $I_{PSI}$ | Slope $I_{PSI}$/[PSI] | Slope $I_{PSII}$/[PSII] | $Ant_{PSI}/Ant_{PSII}$ |
| **658 nm** | 1.96 | 1.01 | 1.72 | 2.29 | 0.75 |
| **732 nm** | 1.70 | 0.88 | 1.49 | 2.00 | 0.75 |

| *C. thermalis* | | | | | |
|---|---|---|---|---|---|
| [PSI]/[PS Tot] | [PSII]/[PS Tot] | PSI/PSII | | Slope $I^{732}/I^{658}$ | |
| 0.84 | 0.16 | 5.13 | | 0.46 | |
| | Slope $I_{PSII+PSI}$ | Slope $I_{PSI}$ | Slope $I_{PSI}$/[PSI] | Slope $I_{PSII}$/[PSII] | $Ant_{PSI}/Ant_{PSII}$ |
| **658 nm** | 2.60 | 2.04 | 2.44 | 3.40 | 0.72 |
| **732 nm** | 1.20 | 1.15 | 1.38 | 0.30 | 4.65 |

Table S1. PSI and PSII antenna size parameters calculated from the maximal electron transport rates from Fig. 7. *I* rates were measured in *S. elongatus*, *A. marina* and *C. thermalis* cells by ECS based on a 200 µs illumination with a 658 nm and a 732 nm actinic LEDs both in the absence and the presence of DCMU and HA. All *I* rates were calculated by normalising on the total amounts of PSI+PSII, measured by flash-dependent ECS. The relative amounts of PSI and PSII ([PSI] and [PSII]) over the total amount of photosystems and the PSI/PSII ratios were calculated based on the amplitudes of flash-dependent ECS measured in the absence and presence of DCMU and HA. Also reported are the slopes of the linear fits of $I_{PSII+PSI}$ and $I_{PSI}$ (measured in the absence and the presence of DCMU and HA, respectively) as a function of light irradiance (in W m$^{-2}$). When measurements were performed in *S. elongatus* with 732 nm illumination, the data could not be fitted with a linear function. The ratio between the PSI and PSII antenna size per reaction centres ($Ant_{PSI}/Ant_{PSII}$) was calculated by normalising the slopes of $I_{PSI}$ and $I_{PSII}$ (the latter calculated as slope $I_{PSII+PSI}$ – slope $I_{PSI}$) on [PSI] and [PSII], respectively, and then taking the ratio between the two values.

| ***S. elongatus*** | | | | | |
|---|---|---|---|---|---|
| **[PSI]/[PS Tot]** | **[PSII]/[PS Tot]** | **PSI/PSII** | | **Slope $I^{732}/I^{658}$** | |
| 0.76 | 0.24 | 3.20 | | 0.18 | |
| | **Slope $I_{PSII+PSI}$** | **Slope $I_{PSI}$** | **Slope $I_{PSI}$/[PSI]** | **Slope $I_{PSII}$/[PSII]** | **$Ant_{PSI}/Ant_{PSII}$** |
| **658 nm** | 1.97 | 1.51 | 1.99 | 1.92 | 1.04 |
| **732 nm** | 0.35 | / | / | / | / |

| ***A. marina*** | | | | | |
|---|---|---|---|---|---|
| **[PSI]/[PS Tot]** | **[PSII]/[PS Tot]** | **PSI/PSII** | | **Slope $I^{732}/I^{658}$** | |
| 0.76 | 0.24 | 3.18 | | 1.36 | |
| | **Slope $I_{PSII+PSI}$** | **Slope $I_{PSI}$** | **Slope $I_{PSI}$/[PSI]** | **Slope $I_{PSII}$/[PSII]** | **$Ant_{PSI}/Ant_{PSII}$** |
| **658 nm** | 1.75 | 1.13 | 1.48 | 2.60 | 0.57 |
| **732 nm** | 2.39 | 1.18 | 1.55 | 5.03 | 0.31 |

| ***C. thermalis*** | | | | | |
|---|---|---|---|---|---|
| **[PSI]/[PS Tot]** | **[PSII]/[PS Tot]** | **PSI/PSII** | | **Slope $I^{732}/I^{658}$** | |
| 0.81 | 0.19 | 4.20 | | 0.73 | |
| | **Slope $I_{PSII+PSI}$** | **Slope $I_{PSI}$** | **Slope $I_{PSI}$/[PSI]** | **Slope $I_{PSII}$/[PSII]** | **$Ant_{PSI}/Ant_{PSII}$** |
| **658 nm** | 0.99 | 0.86 | 1.06 | 0.67 | 1.60 |
| **732 nm** | 0.72 | 0.67 | 0.83 | 0.28 | 3.00 |

Table S2. PSI and PSII antenna size parameters calculated from the maximal electron transport rates from Fig. S7 (independent biological replicates with respect to Fig. 7). *I* rates were measured in *S. elongatus*, *A. marina* and *C. thermalis* cells by ECS based on a 200 µs illumination with a 658 nm and a 732 nm actinic LEDs both in the absence and the presence of DCMU and HA. All *I* rates were calculated by normalising on the total amounts of PSI+PSII, measured by flash-dependent ECS. The relative amounts of PSI and PSII ([PSI] and [PSII]) over the total amount of photosystems and the PSI/PSII ratios were calculated based on the amplitudes of flash-dependent ECS measured in the absence and presence of DCMU and HA, as described in section 3.2 of the main text (see Fig. 6). Also reported are the slopes of the linear fits of $I_{PSII+PSI}$ and $I_{PSI}$ (measured in the absence and the presence of DCMU and HA, respectively) as a function of light irradiance (in W $m^{-2}$). When measurements were performed in *S. elongatus* with 732 nm illumination, the data could not be fitted with a linear function. The ratio between the PSI and PSII antenna size per reaction centres ($Ant_{PSI}/Ant_{PSII}$) was calculated by normalising the slopes of $I_{PSI}$ and $I_{PSII}$ (the latter calculated as slope $I_{PSII+PSI}$ – slope $I_{PSI}$) on [PSI] and [PSII], respectively, and then taking the ratio between the two values.

### References


[1] I. Vass, D. Kirilovsky, A.-L. Etienne, UV-B radiation-induced donor- and acceptor-side modifications of Photosystem II in the cyanobacterium *Synechocystis* sp. PCC 6803, Biochemistry 38 (1999) 12786–12794. https://doi.org/10.1021/bi991094w.

[2] S. Viola, B. Bailleul, J. Yu, P. Nixon, J. Sellés, P. Joliot, F.-A. Wollman, Probing the electric field across thylakoid membranes in cyanobacteria, Proceedings of the National Academy of Sciences 116 (2019) 21900–21906. https://doi.org/10.1073/pnas.1913099116.

[3] S. Viola, J. Sellés, B. Bailleul, P. Joliot, F.A. Wollman, In vivo electron donation from plastocyanin and cytochrome $c_6$ to PSI in *Synechocystis* sp. PCC6803, Biochimica et Biophysica Acta - Bioenergetics 1862 (2021). https://doi.org/10.1016/j.bbabio.2021.148449.

[4] M.Y. Ho, D.M. Niedzwiedzki, C. MacGregor-Chatwin, G. Gerstenecker, C.N. Hunter, R.E. Blankenship, D.A. Bryant, Extensive remodeling of the photosynthetic apparatus alters energy transfer among photosynthetic complexes when cyanobacteria acclimate to far-red light, Biochimica et Biophysica Acta - Bioenergetics 1861 (2020) 148064. https://doi.org/10.1016/j.bbabio.2019.148064.

[5] D.J. Nürnberg, J. Morton, S. Santabarbara, A. Telfer, P. Joliot, L.A. Antonaru, A.V. Ruban, T. Cardona, E. Krausz, A. Boussac, A. Fantuzzi, A.W. Rutherford, Photochemistry beyond the red limit in chlorophyll f–containing photosystems, Science 360 (2018) 1210–1213. https://doi.org/10.1126/science.aar8313.

[6] J. Langley, R. Purchase, S. Viola, A. Fantuzzi, G.A. Davis, J.-R. Shen, A.W. Rutherford, E. Krausz, N. Cox, Simulating the low-temperature, metastable electrochromism of Photosystem I: Applications to *Thermosynechococcus vulcanus* and *Chroococcidiopsis thermalis*, The Journal of Chemical Physics 157 (2022) 125103. https://doi.org/10.1063/5.0100431.

[7] V. Mascoli, A.F. Bhatti, L. Bersanini, H. Van Amerongen, R. Croce, The antenna of far-red absorbing cyanobacteria increases both absorption and quantum efficiency of Photosystem II, Nat Commun 13 (2022) 3562. https://doi.org/10.1038/s41467-022-31099-5.
